# Bacterial and fungal communities are differentially modified by melatonin in agricultural soils under abiotic stress

**DOI:** 10.1101/652388

**Authors:** Andrew P. Madigan, Eleonora Egidi, Frank Bedon, Ashley E. Franks, Kim M. Plummer

## Abstract

An extensive body of evidence from this last decade indicates that melatonin enhances plant resistance to several biotic and abiotic stressors. This has led to an interest in the use of melatonin in agriculture to reduce negative physiological effects from environmental stresses that affect yield and crop quality. However, there are no reports regarding the effects of melatonin on soil microbial communities under abiotic stress, despite the importance of microbes for plant root health and function. Three agricultural soils associated with different land usage histories (pasture, canola or wheat) were placed under abiotic stress by cadmium (100 or 280 mg/kg) or salt (4 or 7 g/kg) and treated with melatonin (0.2 and 4 mg/kg). Automated Ribosomal Intergenic Spacer Analysis (ARISA) was used to generate Operational Taxonomic Units (OTU) for microbial community analysis in each soil. Significant differences in richness (α diversity) and community structures (β diversity) were observed between bacterial and fungal assemblages across all three soils, demonstrating the effect of melatonin on soil microbial communities under abiotic stress. The analysis also indicated that the microbial response to melatonin is governed by the type of soil and history. The effects of melatonin on soil microbes needs to be regarded in potential future agricultural applications.

## Introduction

Soil microbial communities have an essential role in maintaining ecosystem health by direct exchange of nutrients and minerals with plants within the rhizosphere, as well as by providing nourishment to plants indirectly via nutrient cycling of organic matter (Wall and Virginia, 1999, Yao et al., 2000, Kirk et al., 2004, Wahid et al., 2016). Microbial community structures in soil can be altered by various abiotic stresses such as soil contaminants and salinity (Badri and Vivanco, 2009, Wood et al., 2016, Geisseler et al., 2017). Anthropogenic activities, such as phosphorus fertiliser treatment, may introduce the highly toxic heavy metal cadmium (e.g. 300 mg Cd / kg P) into the environment, while irrigation with contaminated water may introduce cadmium or salt (Jiao et al., 2012, Khairy et al., 2014, Roberts, 2014, Bencherif et al., 2015). Both cadmium toxicity and soil salinization cause a dramatic increase in cellular levels of highly destructive reactive oxygen species (ROS) as well as inhibiting the activities of ROS scavenging enzymes in microbes (Tanaka et al., 2006, Achard-Joris et al., 2007, Hossain et al., 2012), thus decreasing crop yield. Globally, up to 2 million hectares of agricultural land are negatively impacted by salinization each year (Bencherif et al., 2015, Ke et al., 2017). Alterations to the soil microbial community can have important repercussions in agricultural settings, where microbes (both soil bacteria and fungi) are critical drivers of soil health and agricultural crop productivity (Mau et al., 2019, Zhang et al., 2019).

Melatonin (N-acetly-5-methoxytryptamine) is an indoleamine (secondary metabolite) produced by all cellular organisms (Hardeland, 2015, Manchester et al., 2015), that can decrease the physiological deleterious effect of abiotic stresses. Melatonin can act either as a highly efficient antioxidant, scavenging up to 10 ROS per molecule, or as a signalling molecule regulating enzymes or hormones associated with ROS scavenging (Weeda et al., 2014, Reiter et al., 2015, Zhang et al., 2015, Hardeland, 2016, Reiter et al., 2016). Melatonin exposure alters gut microbiota composition (Schultz et al., 2006, Chen et al., 2011) and may have antibiotic activity against some microbes (Lucchelli et al., 1997). In plants, exogenous melatonin application via seed-coatings (Wei et al., 2015), soil treatments (Cui et al., 2017), or foliar sprays (Zhang et al., 2017) have been described to promote growth and protect plants against stressors such as cadmium (Byeon et al., 2015, Li et al., 2016, Gu et al., 2017) and salt (Li et al., 2012, Liang et al., 2015, Jiang et al., 2016). However, very little is known about the effects of this secondary metabolites on individual microbes or mixed microbial communities in soil (Tan et al., 2014, Manchester et al., 2015, Paulose et al., 2016).

We examined the effects of melatonin on microbial community structures in three different agricultural soils artificially stressed with cadmium or salt. We hypothesised that melatonin would impact microbial community structures in soils (1) unstressed and (2) under abiotically stressed conditions. We expected specific OTUs to be particularly responsive under the various conditions. This is the first study to date to report significant responses of microbes to melatonin under abiotic stress in agricultural soils at a community level.

## Materials & method

### Soils origin, sampling and physiochemical characteristics

Three agricultural soils associated with different land use were collected from sites within Victoria, Australia. Prior to sampling in late March and early April 2017, site “P” (37°32’28.1’’ S, 145°05’42.5‘’ E) was most recently (< 3 months) associated with cattle and sheep <u>P</u>asture, site “C” (37°45′28.4″ S, 144°14′30.2″ E) was 3 weeks post <u>C</u>anola harvest, and site “W” (37°43′31.1″ S, 144°13′14.3″ E) was 3 weeks post fire blazing of stubble following a <u>W</u>heat harvest. As the plant species associated with a soil has been shown to be a key factor influencing the bacterial community structure within the rhizosphere (Burns et al., 2015), it was expected that selecting these soils would provide the most diverse range of soil microbes for the current study.

At each site, approximately 4 kg topsoil was sampled (10 cm deep × 0.5 m width × 0.5 length) from each of four plots spaced 3 m apart. Soils were air dried overnight and sieved to remove particles larger than 2 mm. A single soil sample was generated for each site by pooling all four collected soils from the subsite plots (Aye et al., 2016, Butterly et al., 2016). The collected stock soils were separately stored at ambient room temperature (21°C) in airtight plastic containers for four months, followed by subsampling (representing the first sampling timepoint; labelled as ‘T0’ in the analysis) for immediate treatments as described below. During this first soil subsampling process stock soils were thoroughly aerated. The stock soils were then stored at ambient room temperature (21°C) in airtight plastic containers for a further four weeks and subsampled for immediate treatments (representing the second sampling timepoint; labelled as ‘T1’ in the analysis). It was expected that this aging regime would change the initial microbial populations and potentially shift selection pressures amongst the populations (Kelly et al., 1999, Castro et al., 2010, Heijboer et al., 2018, Reese et al., 2018). Physicochemical analyses of untreated soils were conducted by Nutrient Advantage (Melbourne, Australia) (Table 1). Electrical conductivity (EC) of untreated and salt-treated (NaCl) soils was determined using 1:5 soil:water as per He *et al*. (He et al., 2012) (Table 2).

**Table 1:** Physical and chemical characteristics of the three agricultural soils used int his study. Four topsoil samples (0-10 cm) were collected at each site and composited prior to analysis.

| Site | P | C | W |
| --- | --- | --- | --- |
| Available Potassium (mg kg<sup>-1</sup> soil) | 640 | 140 | 170 |
| pH (1:5 CaCl<sub>2</sub>) | 6.1 | 4.9 | 4.7 |
| Organic Carbon (%) | 4.6 | 3 | 2.8 |
| Nitrate N (mg kg<sup>-1</sup> soil) | 160 | 57 | 34 |
| Ammonium N (mg kg<sup>-1</sup> soil) | 2.5 | 8.9 | 8.8 |
| Phosphorus (Colwell) (mg kg<sup>-1</sup> soil) | 160 | 41 | 32 |
| Phosphorus Buffer Index | 32 | 71 | 83 |
| Calcium (cmol(+) kg<sup>-1</sup> soil) | 12 | 5.1 | 4.7 |
| CEC (cmol(+) kg<sup>-1</sup> soil) | 15.7 | 7.94 | 8.56 |
| EC (1:5 water) (dS m<sup>-1</sup>) | 0.37 | 0.24 | 0.18 |
| Chloride (mg kg<sup>-1</sup> soil) | 37 | 67 | 76 |
| Moisture (%) | 14 | 6 | 5.2 |
| Water holding capacity (%) | 38.4 | 24.6 | 21.1 |
| Sand (%) | 75.2 | 53.9 | 52.4 |
| Silt (%) | 23.6 | 29.9 | 28.8 |
| Clay (%) | 1.2 | 16.2 | 18.8 |
| Iron (mg kg<sup>-1</sup> soil) | 5400 | 16000 | 21000 |
| Zinc (mg kg<sup>-1</sup> soil) | 23 | 7.2 | 9.3 |
| Cadmium (mg kg<sup>-1</sup> soil) | 0.24 | 0.13 | 0.2 |
| Chromium (mg kg<sup>-1</sup> soil) | 11 | 17 | 32 |
| Nickel (mg kg<sup>-1</sup> soil) | 4 | 4.7 | 22 |
| Lead (mg kg<sup>-1</sup> soil) | 7.7 | 9.4 | 16 |

CEC = cation exchange capacity; EC = electrical conductivity; Soil P – recently (< 3 months) associated with cattle and sheep pasture; Soil C – sampled 3 weeks post canola harvest; Soil W –sampled 3 weeks post fire blazing following a wheat harvest.

**Table 2:** Electrical conductivity (milli Siemens / centimetre) [mS / cm] in soils P, C and W treated with high (7 g kg^−1^ soil) and low (4 g kg^−1^ soil) salt (NaCl). Control treatments consist of sterile Milli-Q water replacing salt solutions. All treatments and controls within the same soil were significantly different (p < 0.05) from each other as determined by one-way ANOVA. Salinity levels as measured by Electrical conductivity [EC (1:5 soil:water)]. SE: Standard error

| EC 1:5 (mS/cm) |  |  |  |  |  |  |  |  |  |
| --- | --- | --- | --- | --- | --- | --- | --- | --- | --- |
|  | Soil P |  |  | Soil C |  |  | Soil W |  |  |
|  | 7 g kg<sup>-1</sup> | 4 g kg<sup>-1</sup> | Control | 7 g kg<sup>-1</sup> | 4 g kg<sup>-1</sup> | Control | 7 g kg<sup>-1</sup> | 4 g kg<sup>-1</sup> | Control |
| Rep 1 | 2.6 | 1.72 | 0.706 | 1.97 | 1.015 | 0.091 | 1.934 | 1.148 | 0.101 |
| Rep 2 | 2.545 | 1.708 | 0.678 | 1.943 | 1.083 | 0.132 | 1.954 | 1.089 | 0.093 |
| Rep 3 | 2.502 | 1.8 | 0.681 | 1.829 | 1.084 | 0.138 | 1.983 | 1.065 | 0.102 |
| **Mean** | **2.549** | **1.743** | **0.688** | **1.914** | **1.061** | **0.120** | **1.957** | **1.101** | **0.099** |
| SE | 0.028 | 0.029 | 0.009 | 0.043 | 0.023 | 0.015 | 0.014 | 0.025 | 0.003 |

### Soils treatment with abiotic stress (cadmium or salt) and melatonin

The responses of soil microbial communities to exogenous melatonin application was investigated with and without cadmium and salt as separately applied stressors (all chemicals from Sigma Aldrich Pty. Ltd., Australia). Five grams of sieved topsoil was transferred to a sterile 50 ml Falcon tube and exposed to various treatment combinations. At sampling timepoint T0, treatments were composed of high or low melatonin (4 or 0.2 mg kg^−1^ soil respectively) and / or high or low cadmium (cadmium chloride hemipentahydrate) (280 or 100 mg kg^−1^ soil respectively). At sampling timepoint T1, dry (untreated) soils were treated with a solution composed of melatonin (4 or 0.2 mg kg^−1^ soil) and / or high or low salt (NaCl) stressor (7 or 4 g kg^−1^ soil respectively). Controls involved dilute ethanol replacing melatonin (see below) and/or sterile Milli-Q water replacing cadmium. These concentrations were selected to be within the range of those reported to induce effects on soil microbial activities in previous studies for cadmium (Cáliz et al., 2013, Wood et al., 2016) and salt (Rath et al., 2016). Based upon preliminary studies, soils were treated to 80-90% field capacity, to ensure all soils were sufficiently exposed to stressor and melatonin. Soils without the addition of a stressor acted as a control.

Melatonin was initially dissolved in 99.9% ethanol to 200 mM and diluted to the required concentrations with sterile Milli-Q water. All treatments and controls contained a standardised amount of ethanol (100 μl of 0.43% ethanol per 5 g soil). Due to differing water holding capacities for each soil, this equated to a final ethanol concentration of 0.044%, 0.052% and 0.06% v/v in treatments within soils P, C and W respectively. Control treatments were composed of dilute ethanol replacing melatonin and sterile Milli-Q water replacing cadmium (sampling timepoint T0) or NaCl (sampling timepoint T1). Treatments and controls were conducted in quadruplicates. Samples were incubated in sterile 50 mL falcon tubes covered with loosely fitted lids to enable gas exchange at room temperature (21°C) in darkness for 10 days. Four untreated samples (i.e. no water added) replicates per soil were also collected on Day 0 to provide a representation of the baseline communities prior to treatments.

### DNA extraction and amplification for Automated Ribosomal Intergenic Spacer Analysis (ARISA)

Automated Ribosomal Intergenic Spacer Analysis (ARISA) is a molecular technique used to characterise the microbial diversity within various environments, including bulk soil and the rhizosphere (Sørensen et al., 2009, Rincon-Florez et al., 2013). This technique fingerprints fungal and bacterial communities based upon the length heterogeneity of the Intergenic spacer region between the ribosomal Ribonucleic Acid (rRNA) genes for bacteria (16S and 23S) and/or fungi (18S and 28S) (Kirk et al., 2004). ARISA provided a valuable estimate of both species richness and relative abundances, allowing a comparison of *α* and *β* diversity both across various sites and between samples exposed to different treatments (Zancarini et al., 2012).

Total soil DNA was extracted from 0.3 g of soil subsamples using PowerSoil^®^ DNA Isolation Kit (MoBio Laboratories Inc., California, U.S.A.) according to manufacturer’s instructions. For the fungal community analysis, the internal transcribed spacer regions 1 and 2 (ITS 1 and 2) were amplified using fungal primers ITS-1F and ITS-4 (Blaalid et al., 2013). A 20 μl fungal ARISA PCR mastermix contained 1 × PCR buffer (Qiagen Pty Ltd., Melbourne, Australia); 1.5 mM MgCl_2_ (Qiagen); 1 × Q reagent (Qiagen); 500 μM concentration of each deoxyneucleoside triphosphate (dNTP) (Qiagen); 10 ng of extracted DNA; 500 nM of fungal primers ITS 1F (5′-CTT GGT CAT TTA GAG GAA GTA-3′) and ITS 4 (5′-TCC TCC GCT TAT TGA TAT GC-3′) (Bioline Global Pty Ltd., NSW, Australia), the latter labelled with a phosphoramidite dye, 6FAM (Sigma Aldrich Pty Ltd, Sydney, Australia); and 3.75U of GoTaq polymerase (Qiagen). Reaction mixtures were held at 96°C for 3 min, followed by 36 cycles of amplification at 96°C for 30 sec, 55°C for 75 sec and 72°C for 90 sec and a final extension of 72°C for 6 min.

Bacterial community analysis by ARISA targeted the intergenic spacer region of the 16S – 23S rRNA genes using the universal primer 16S – 1392f (5′ - GSA CAC ACC GCC CGT - 3′), labelled with a phosphoramidite dye, 6FAM, and bacterial primer 23S – 125r (5′ - GGG TTB CCC CAT TCR G - 3′) (Fuhrman et al., 2008). The bacterial ARISA PCR mixture contained the same ingredients and concentrations as applied in the fungal PCR mastermix (described above), with the exception of the primers used. Reaction mixtures were held at 94°C for 3 min, followed by 33 cycles of amplification at 94°C for 1 min, 52°C for 1 min and 72°C for 90 sec and a final extension of 72°C for 6 min. PCR products were examined by gel electrophoresis on a 1.5% agarose gel matrix and imaged by fluorescence under UV light. Bacterial and fungal community analyses by ARISA were conducted by Australian Genome Research Facility as per Wood et al. (2016) (Wood et al., 2016). This involved the separation of amplicons, described as operation taxonomic units (OTUs), by capillary electrophoresis.

### Statistical Analysis of ARISA data

OTU fragment sizes were limited to a range of between 140 – 1000 bp for both fungi and bacteria to ensure only the intergenic spacer region was represented in the data (Fisher and Triplett, 1999). Singletons and low abundance amplicons (<1% relative abundance) were disregarded (Fisher and Triplett, 1999). Data were normalized in the statistical software R version 3.3.2 (The R Foundation for Statistical Computing, Boston, USA), with bin sizes of 3 and 4 selected for bacteria and fungi respectively (Ramette, 2009, Butterly et al., 2016). The following analyses were conducted in Primer-E v6 (Quest Research Ltd., Auckland, New Zealand), with treatments considered as fixed factors and microbial responses analysed for each soil separately (Chow et al., 2013, Wood et al., 2016): 1) Bray-Curtis similarity for microbial communities; 2) SIMPER analysis to identify the contribution of individual Operational Taxonomic Units (OTUs) to (dis)similarity between replicates or different treatments; 3) DIVERSE to determine Shannon diversity index (H’) for sample data; 4) Permutational multivariate analysis of variance (PERMANOVA) to determine treatment effect (melatonin, stressors: cadmium and Salt) on microbial assemblages for each soil at various concentrations (High, Low and Zero). Monte Carlo statistical analysis was conducted to determine if individual treatment combinations had statistically significant effects on community compositions (Van Wijngaarden et al., 1995). Shannon’s diversity index (H’) was calculated from binned data to determine differences in biodiversity (relative abundance and evenness) of taxa present within each soil for fungal and bacterial communities upon treatment with melatonin (averaged for 4 replicates) (Ondreičková et al., 2016). Subsequently, Shannon index values were analysed by non-parametric Wilcoxon test to determine significant (p < 0.05) differences upon melatonin treatments under high or low concentration for the stressors (cadmium or Salt). Non-multidimensional scaling (nMDS) and principal coordinates Analyses (PCoA) plots were generated using Analysis of Similarity (ANOSIM) default parameters (significance [p < 0.05], 9999 permutations) for visual representation of community (dis)similarities between samples.

### Microbial biomass response to melatonin and stressors

Bacterial and fungal biomasses were measured by quantitative PCR (qPCR) on DNA of the 16s rRNA and ITS region respectively. Bacterial communities were assessed using primer pairs 1114f (5’-CGG CAA CGA GCG CAA CCC-3’) −1275r (5’-CCA TTG TAG CAC GTG TGT AGC C-3’), and fungal communities were assessed using ITS1F (5’-TCC GTA GGT GAA CCT GCG G-3’) −5.8Sr (5’-CGC TGC GTT CTT CAT CG-3’) primer (Wood et al., 2016). Each treatment consisted of four biological replicates each quantified in technical triplicate using a CFX Connect Real-Time PCR Detection System (BioRad). A 20 μl reaction mixture for bacterial samples was composed of 2 μl extracted DNA (0.25 ng/μl) and 18 μl mastermix according to the following recipe: 3.3 μl Universal SYBR^®^ Green Super Mix; 0.27 μl of each 10 μM forward and reverse primer (135 pM final concentration of each primer in reaction mixture); 14.16 μl DNA-free water. Bacterial qPCR reaction mixtures were held at 94°C for 3 min, followed by 40 cycles of amplification at 94°C for 10 sec, 61.5°C for 30 sec. Fungal samples were prepared to a 10 μl reaction mixture composed of 2 μl extracted DNA (0.25 ng/μl) and 8 μl mastermix composed of 4.5 μl Universal SYBR^®^ Green Super Mix; 0.5 μl of each 10 μM forward and reverse primer (500 pM final concentration of each primer in reaction mixture); 2.5 μl DNA-free water. Fungal qPCR conditions were 95°C for 5 min, followed by 40 cycles of amplification at 95°C for 30 sec, 53°C for 30 sec, and 72°C for 30 sec. A melting curve was measured from 65°C up to 95°C following qPCR reactions by increasing in 0.5°C increments every 30 sec. Purified amplicons from pure isolates of *E. coli* and *Penicillium* sp. cultures were used to generate standard curves (10-fold series dilutions) for bacterial and fungal samples respectively.

## Results

### Agricultural soils have diverse microbiomes and physiochemical properties

Three agricultural soils differing in land use practices, physiochemical characteristics and microbial content were used to assess the modulatory effect of melatonin on soil community. Soils C and W were collected from sites within close geographical proximity (< 10 km) to each other and showed the most similarity in physiochemical parameters of the three soils (Table 1). Electrical conductance significantly increased in response to both salt treatments treatment (Table 2). The ARISA community profiles representing bacterial and fungal communities in the three dry untreated soils showed significant (p < 0.05) differences to each other (Figure 1). We detected a mean of 29.7, 30 and 36 bacterial OTUs, and 16.6, 16.4 and 14.8 fungal OTUs, for untreated soils P, C and W respectively.

**Figure 1:**
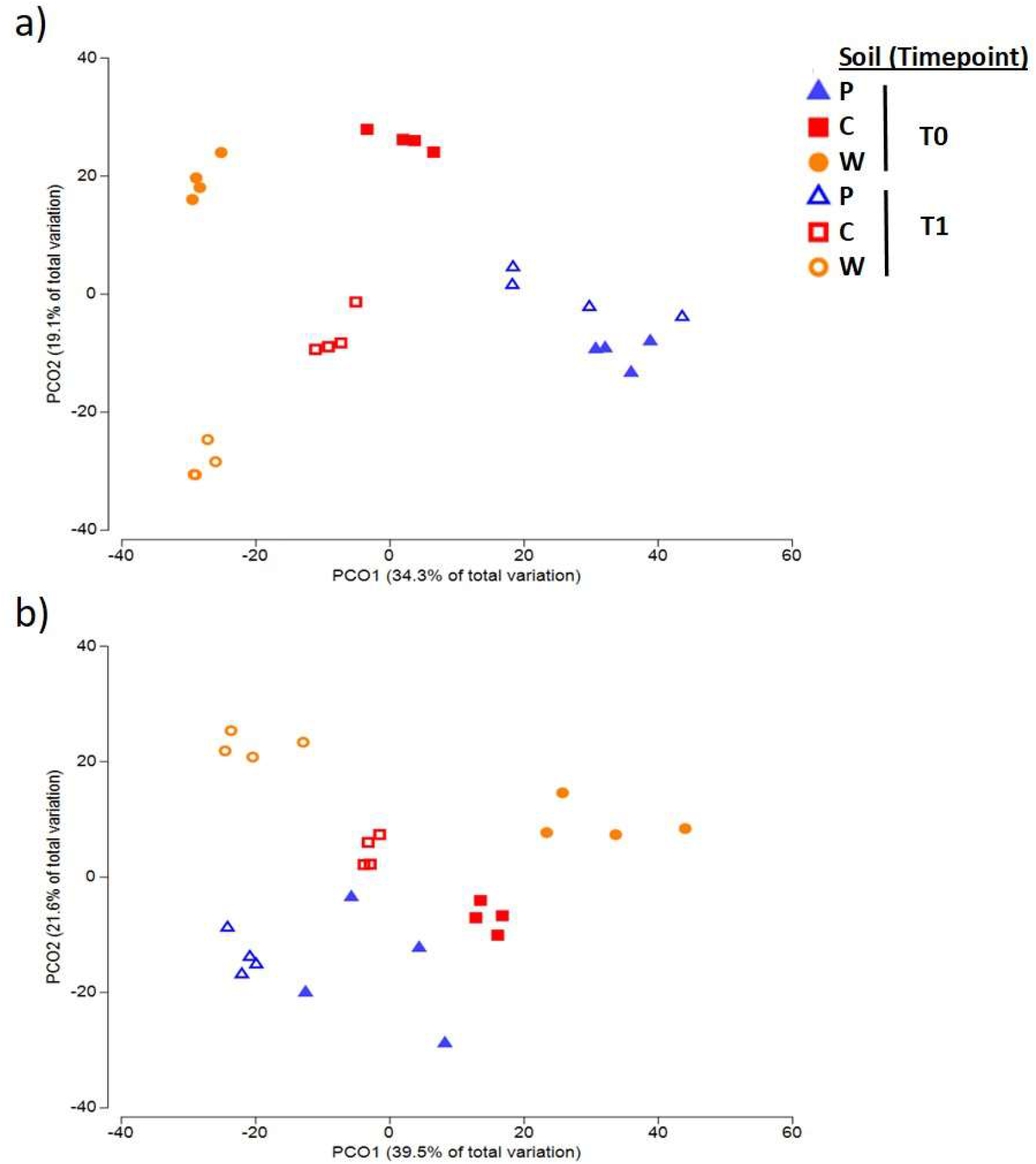
Principal coordinates analysis (PCOA) ordination plots (Bray-Curtis distance matrix) of ARISA profiles showing the separation within a) bacterial and b) fungal communities for the three dry, untreated soils (P, C, W; n = 4). Bacterial communities from the same soil differed (ANOSIM: R=0.865 − 1; p = 0.029) between sampling timepoints T0 and T1 within soils C and W, whereas bacterial communities in soil A were not significantly different between timepoints T0 and T1 (ANOSIM: R=0.594; p = 0.057). All untreated soil communities from different soils were dissimilar (p < 0.05) to each other. Fungal communities from the same soil differed (ANOSIM: R = 0.458 − 1; p = 0.029) between T0 and T1 for all three soils.

### Cadmium and salt alter microbial community structures

Stressors significantly (p < 0.05) impacted microbial β diversity (Table 3). Bacterial community structures of soils C and W were significantly (*p* < 0.01) impacted by treatments with high and low concentrations of cadmium and salt (Supplementary Table 1a). Similarly, bacterial community structures in soil P were responsive to salt at high and low concentrations, but community structures in this soil did not show a significantly response to cadmium. Fungal communities in all soils were impacted by high concentrations of cadmium and salt. In contrast, low concentrations of salt impacted fungal community structures in soil C and W (not P), whereas only fungal communities in soil P were altered by low cadmium treatment (Supplementary Table 1b).

**Table 3:**
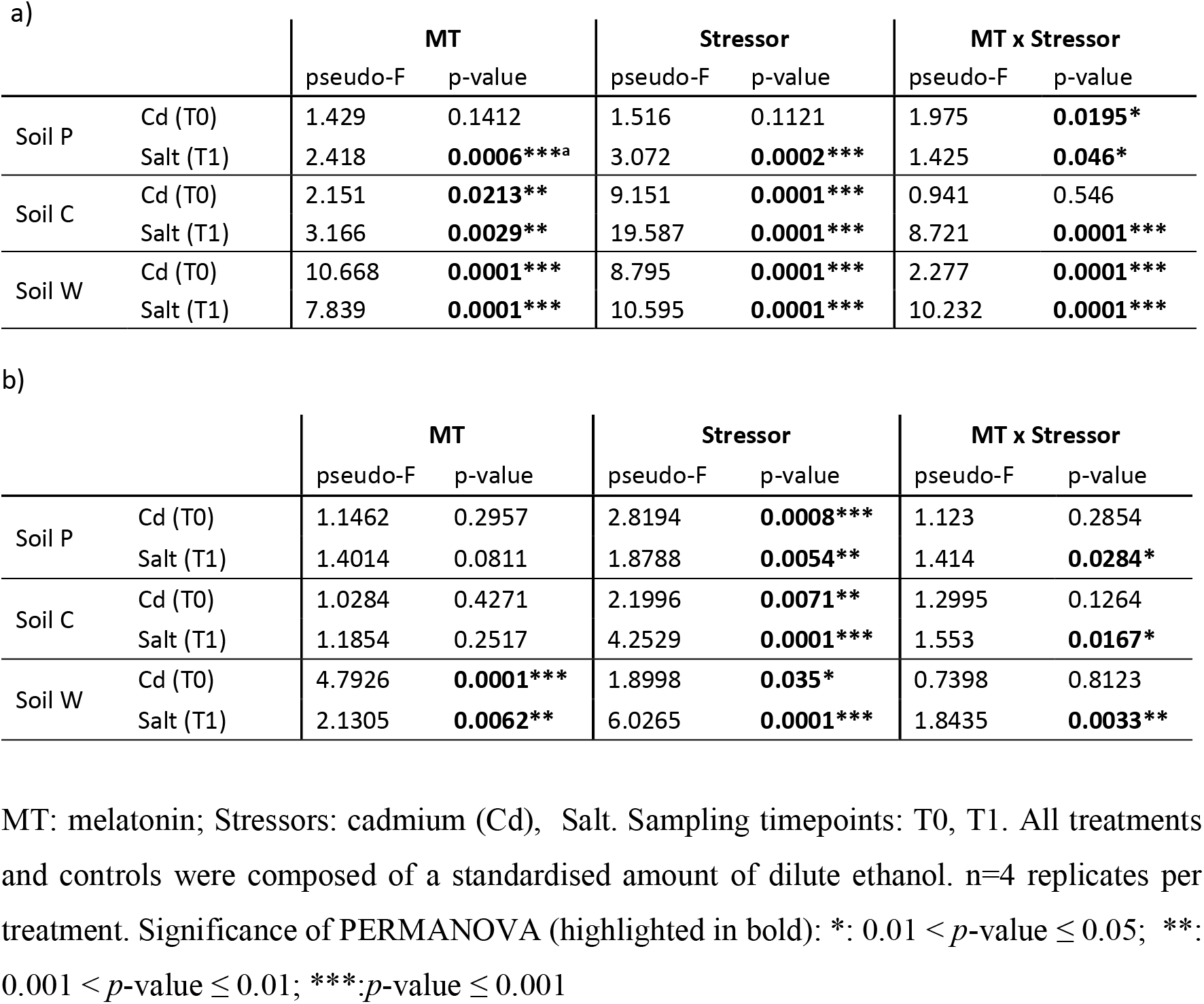
Bacterial a) and fungal b) community responses to treatments (β diversity) with melatonin, stressors and melatonin-stressor combinations based on PERMANOVA analyses of ARISA data (Bray-Curtis dissimilarity distances).

### Melatonin affects soil microbial diversity and abundance across all three soils

#### a) Soil microbial community responses to melatonin under abiotic stress conditions

The addition of melatonin to all three soil types had significant effects (increases and decreases) on the bacterial α diversity (*p* < 0.05) but did not significantly affect the fungal α diversity (Supplementary Data 1 [Supplementary Table 2]). Individual bacterial and fungal OTUs that responded strongly to melatonin showed relative abundances increased by up to 7.1% and 11.5% and reduced by 16.9% and 10.2% respectively (Supplementary Tables 3 & 4). Analysis of the soils OTUs compositions (β diversity) by Permutational Multivariate Analysis of Variance (PERMANOVA) indicated that bacterial and fungal community were significantly altered by melatonin and stressors application (Table 3, Supplementary Tables 1, 5 & 6). The responses of microbial communities in all soils to melatonin under abiotic stress conditions was visualised by non-metric multidimensional scaling (nMDS) ordination (Figures 2a & 3a; Supplementary Figures 1 & 2). Under abiotic stress conditions, bacterial communities responding significantly (*p* < 0.05) to melatonin showed a decreased Shannon diversity index (OTU abundance and evenness), whereas fungal communities increased under the same conditions (Supplementary Data 2 [Supplementary Figures 3 & 4]). Overall melatonin had very little effect on bacterial 16S or fungal 18S rDNA copy numbers, with only one soil bacterial community (Soil W) significantly impacted (*p* < 0.05) by exogenous melatonin only, while fungal communities were unaffected by melatonin-only treatments (Supplementary Data 3 [Supplementary Figures 5 & 6]). Some significant differences between treatments were recorded, however no consistent pattern of shift in microbial biomass under stress conditions +/− melatonin was observed (Supplementary Data 3).

**Figure 2:**
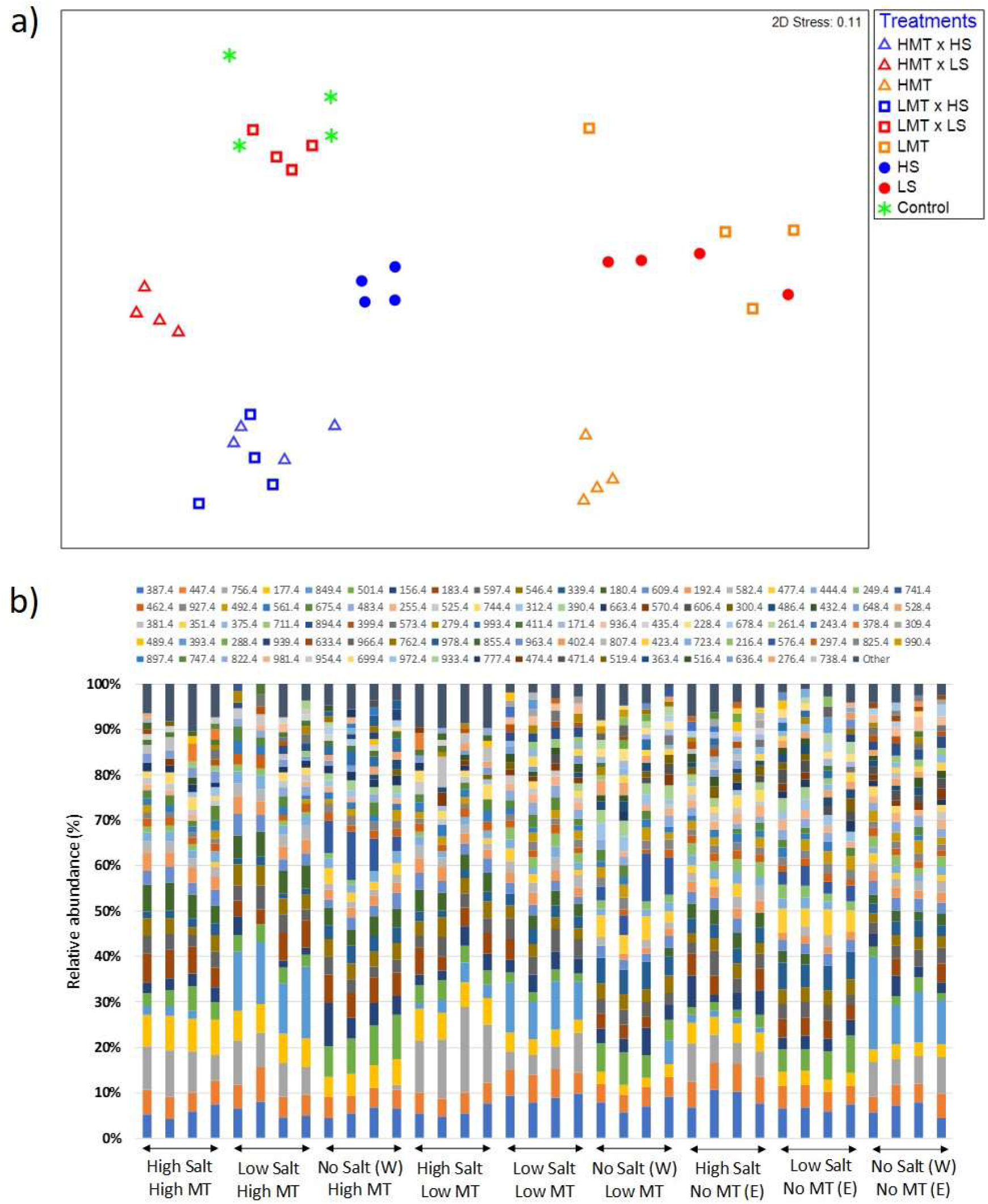
Comparison of a) non-metric multidimensional scaling (nMDS) ordination and b) OTU relative abundances displaying Bray-Curtis similarities for bacterial samples within soil W for various treatments of melatonin and salt based upon community compositions determined by ARISA fingerprinting analysis. Low 2D stress value (< 0.20) indicate high quality ordination plots. Legends in figure 2b) represent different individual OTUs as determined by specified nucleotide lengths. Relative proximity of replicates reflects high community similarity within the same treatments for bacterial communities. Water (W) replaced salt treatment and dilute ethanol (E) replaced melatonin treatments in respective control samples. All treatments and controls were composed of a standardised amount of dilute ethanol. (HMT: High melatonin; LMT: Low melatonin; HS: High salt; LS: Low salt; HMT × HS: High melatonin with high salt; HMT × LS: High melatonin with low salt; LMT × HS: low melatonin with high salt; LMT × LS: Low melatonin with low salt).

**Figure 3:**
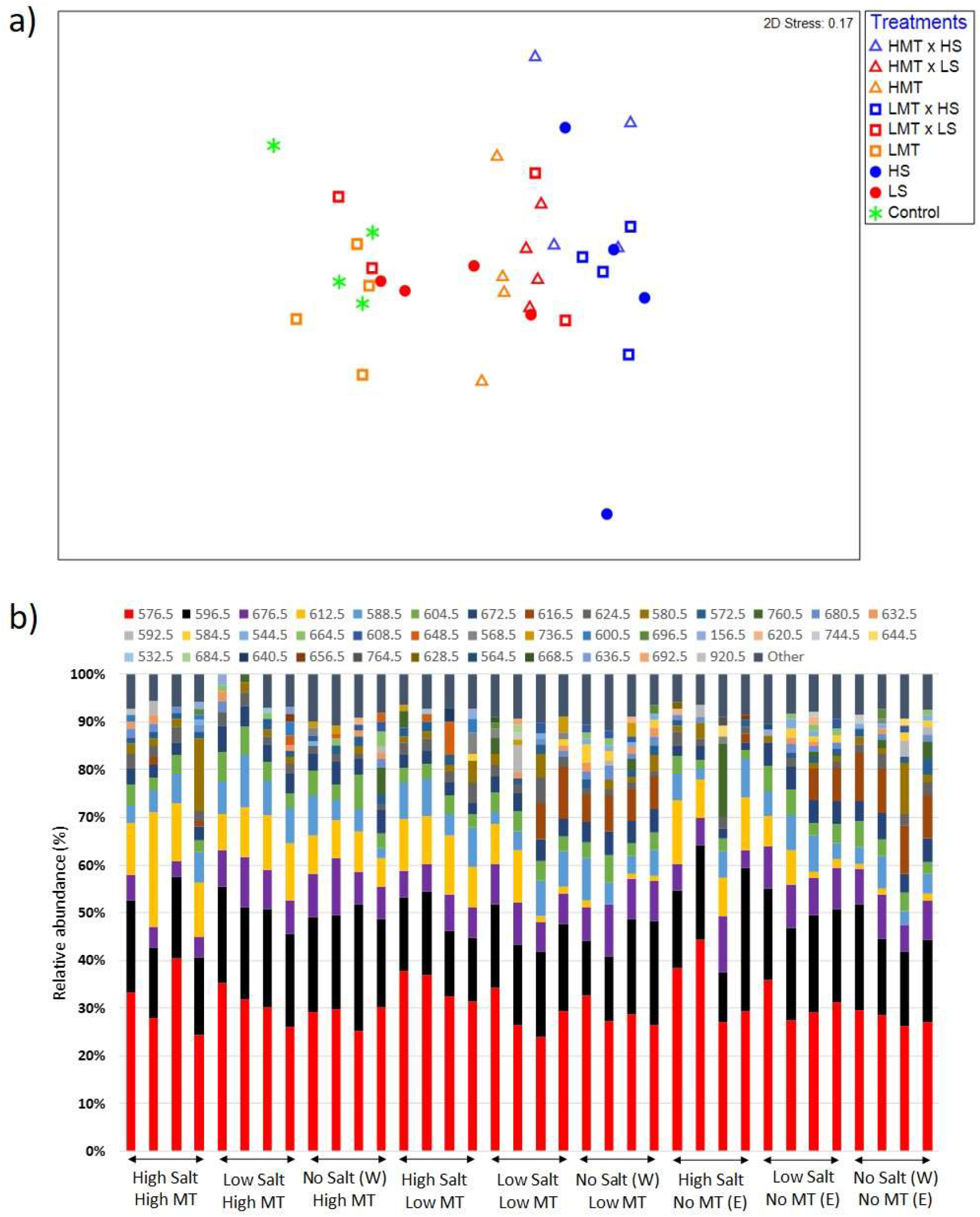
Comparison of a) non-metric multidimensional scaling (nMDS) ordination and b) OTU relative abundances displaying Bray-Curtis similarities for fungal samples within soil W for various treatments of melatonin and salt based upon community compositions determined by ARISA fingerprinting analysis. Low 2D stress value (< 0.20) indicate high quality ordination plots. Legends in figure 3b) represent different individual OTUs as determined by specified nucleotide lengths. Relative proximity of replicates reflects high community similarity within the same treatments for fungal communities. Water (W) replaced salt treatment and dilute ethanol (E) replaced melatonin treatments in respective control samples. All treatments and controls were composed of a standardised amount of dilute ethanol. (HMT: High melatonin; LMT: Low melatonin; HS: High salt; LS: Low salt; HMT × HS: High melatonin with high salt; HMT × LS: High melatonin with low salt; LMT × HS: low melatonin with high salt; LMT × LS: Low melatonin with low salt).

#### b) Effect of melatonin on bacterial and fungal communities

Melatonin had different impacts on bacterial versus fungal composition of soils. High melatonin concentration had a significant effect (PERMANOVA: *p* < 0.01) on bacterial community structures in all three soils, whereas low melatonin concentration impacted only bacterial communities within soil C and W (Supplementary Table 5a). Fungi responded less to melatonin compared to bacteria within the same samples. Melatonin had no effect on fungal community structures in soil C (Supplementary Table 5b), while only high melatonin concentration treatment at sampling timepoint T1 impacted fungal community structures in soil P (Table 3). However, melatonin at high and low concentration induced shifts in fungal community structures within soil W.

#### c) Microbial community responses to melatonin under stress condition

In comparison to stressor-only treatments, consistent responses to melatonin were observed in bacterial communities compared to fungal communities, especially at high melatonin concentration (Table 4; Supplementary Table 6). Bacterial responses to high melatonin concentration were significant (in comparison with stressor only treated communities) under high cadmium or salt conditions in soils P and W, but not in soil C (Table 4). Under low stressor conditions, bacterial communities in soil C and soil W responded significantly to high melatonin treatment. In contrast, bacterial communities responded to the availability of low melatonin under high stress conditions only in soil W only (both stressors) and under low stressor (salt only) in soils C and W. Fungal communities were far less responsive to melatonin treatments under abiotic stress conditions. In comparison with stressor only treatments, melatonin only caused a shift in fungal communities at high melatonin treatments in soil W under high cadmium stress, and to low melatonin under low salt stress in soil C (Table 4).

**Table 4:**
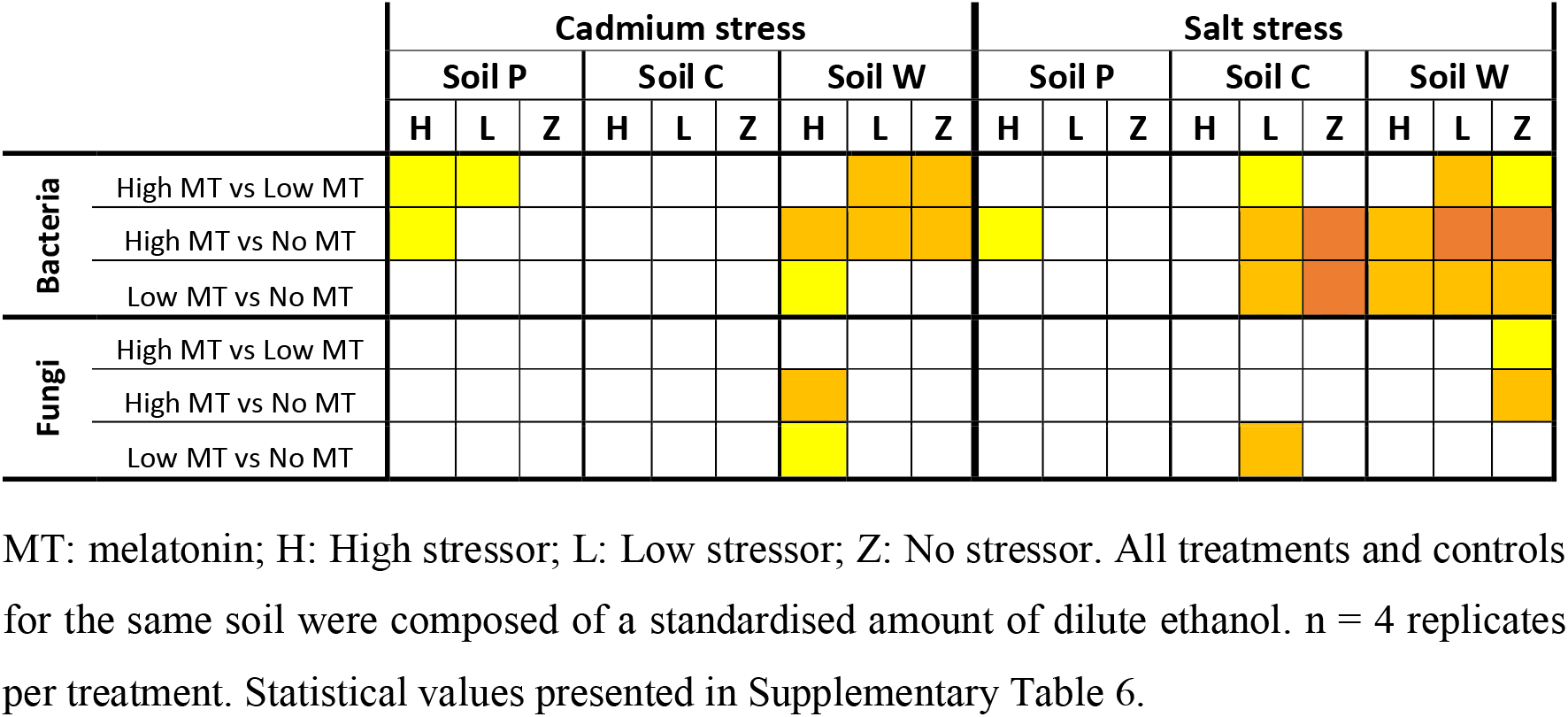
Responses of bacterial and fungal communities to individual treatments of melatonin under various stressor conditions as determined by PERMANOVA. For control treatments, melatonin was replaced with dilute ethanol. Significant differences (p < 0.05) between melatonin treatments are highlighted under the following significance levels: 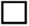 : *p* > 0.05 (n.s.); 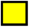: 0.01 < *p* ≤ 0.05; 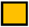: 0.001 < *p* ≤ 0.01; 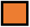: *p* ≤ 0.001

### Microbial responses within soil W communities

Only soil W under salt stress showed significant (*p* < 0.01) responses to treatments of melatonin, stressor and melatonin x stressor for both fungal and bacterial communities as described by PERMANOVA (Table 3; Supplementary Tables 1, 5 & 6). These communities were subsequently assessed in greater detail.

#### a) Bacterial communities show distinct responses to melatonin under salt stress (soil W)

The nMDS plot for bacterial assemblages showed clear separation of samples according to treatments, with both melatonin and salt treatments resulting in shifts in community structures compared to the control (Figure 2a). Low 2D stress values (< 0.20) indicated high quality ordination plots. Various individual OTUs shifted in response to the different treatments (Figure 2b, Supplementary Figure 7a). Bacterial communities treated with melatonin were significantly (PERMANOVA F_(2, 27)_ = 7.839, *p* < 0.001) different to control communities, with high melatonin concentration being 50.5% dissimilar to the control treatments whereas low melatonin concentration samples showed 48.0% dissimilarity (Supplementary Table 7a). Similarity Percentage Analysis (SIMPER) showed the OTUs contributing most to assemblage differences upon high melatonin treatment in comparison to control samples. This analysis found that the top three OTUs accounted for 27% of the total dissimilarity. These were OTU: 849 bp (12.9%), 741 bp (7.3%) and 756 bp (6.8%). High and low concentration melatonin samples were 36.7% dissimilar to each other with 29 OTUs accounting for 71.1% of these differences, the highest of which (OTU: 180 bp) represented only 5.0% of the dissimilarity (Supplementary Figure 7a). Bacterial responses to both concentrations of melatonin for soil W were significant under high and low salt stress in comparison to the respective salt only treatments (Table 4). Under all the above comparisons, a single OTU (756 bp) consistently increased (up to 7.7%) when melatonin was present, independent of melatonin and salt concentrations. Interestingly, some OTUs most responsive to melatonin under high salt conditions in soil W (e.g. OTU 387 bp) were far less responsive to melatonin under low salt conditions and vice versa (e.g. OTU 849 bp).

#### b) Fungal community responses to treatments (melatonin and/or salt) in soil W

NMDS ordination for fungal communities within soil W indicated significant separation of melatonin (PERMANOVA: F_(2, 27)_ = 2.131, *p* < 0.001) and salt (PERMANOVA: F_(2, 27)_ = 6.027, *p* < 0.001) treated samples with respective control samples (Figure 3a & b). Relative abundances of individual fungal OTUs varied across all treatments in soil W under salt stress (Supplementary Figure 7b). Fungal communities in soil W treated with high melatonin concentration showed greater dissimilarity (33.7%) than low melatonin treated samples (28.5%) when both were compared with control samples (Supplementary Table 7b), with high melatonin treatment significantly different (*p* < 0.05) to control samples (Table 4). In soil W communities treated with high melatonin concentration, three taxa accounted for almost half of the total dissimilarity: OTUs 607 bp (18.6%), 575 bp (16.8%) and 591 bp (13.2%). Correspondingly, OTUs 575 bp (19.2%) and 591 bp (13.8%) also accounted strongly for dissimilarity between control and low melatonin-treated samples (Supplementary Figure 7b). Separation between samples treated with low melatonin concentration only compared with samples treated with both salt and low melatonin concentration was observed, however fungal community differences were not determined as significant in response to melatonin under salt stress (Figure 3a; Table 4).

## Discussion

Soil microbial communities are a key component of a healthy ecosystem, providing a number of ecosystem services including direct and indirect nourishments of plant root systems (Wall and Virginia, 1999, Yao et al., 2000, Kirk et al., 2004, Wahid et al., 2016). We detected significant responses of microbes at a community level to melatonin under normal or abiotic stress conditions in agricultural top-soils (10cm depth). Shifts in soil microbial communities may result in changes to various ecosystem services provided by soil microbes (Mau et al., 2019, Zhang et al., 2019). Our data therefore suggests that any potential applications of melatonin in future agricultural practices must also investigate the resulting shift to soil microbial communities to ensure that plant-soil ecology interactions are not negatively impinged in the long term.

We hypothesised that melatonin would alter soil microbial community structures, as previous reports suggest that melatonin may act with antimicrobial properties. For example, melatonin was shown to inhibit the in *vitro* growth of the human bacterial pathogens, *Staphylococcus aureus, Pseudomonas aeruginosa* and *Acinetobacter baumannii* at concentrations between 130-530 μM (Tekbas et al., 2008). Melatonin has also been reported to inhibit the *in vitro* growth of the human pathogenic yeast, *Candida albicans*, albeit at a much higher concentration (1300 μM) (Öztürk et al., 2000). In our study, bacterial communities in three different agricultural soils were affected by melatonin alone, however, fungal communities were less responsive to melatonin compared with bacteria (Table 2; Supplementary Table 6). *In vitro* studies investigating responses of filamentous fungi to melatonin complement our finding as melatonin, at very high concentration (100 mM), showed no impact on the *in-vitro* growth of *Physalospora piricola, Botrytis cinerea* or *Mycosphaerella arachidicola* (Wang et al., 2001). Growth of *Alternaria* spp. has been reported to be inhibited at a 4 mM (Arnao and Hernández-Ruiz, 2015). Taken together our results from soil bacteria analysis indicate a similar trend compared to the *in vitro* experiments albeit different melatonin concentration ranges.

Recently Li et al., (2018) investigated the effect of exogenous melatonin application (200 μM; applied at 20 day intervals for 6 months) without abiotic stress in two soils types associated with horticultural practices (apple orchard and vegetables respectively) by sampling the subsoil region (20-30 cm depth). Bacterial compositions of melatonin-treated soil samples were shown to be similar to controls, however some genera, many unknown, shifted strongly in response to melatonin (Li et al., 2018). Ascomycota, in particular, were negatively affected by melatonin, resulting in greater establishment of *Glomeromycota* and *Basidiomycota* in fungal assemblages (Li et al., 2018). It is difficult to directly compare our study with this report, however, these results are complimentary to the trends observed in our investigation and indicative of the importance of soil agricultural histories in microbial response to melatonin. In our study, we used topsoils, which are generally more microbially rich (compared to subsoils), as well as soils associated with agricultural practices (crop production and pasture). Further analysis by taxa identification of the soil microbial communities associated with responses to melatonin would determine those microbes associated with particular ecological niches (e.g. mycorrhizal fungi or Plant Growth-Promoting Rhizobacteria) and if they are known to be beneficial to crops.

Under abiotic stresses, such as cadmium and salt, plants can cope better by adjusting physiological and enzymatic processes when melatonin is applied (Li et al., 2012, Byeon et al., 2015, Liang et al., 2015, Jiang et al., 2016, Li et al., 2016, Gu et al., 2017). Some reports also suggested that melatonin may enhance abiotic stress tolerance in microbes as endogenous levels of melatonin increased in *Trichoderma* spp. (Liu et al., 2016) and *Saccharomyces cerevisiae* (Rodriguez-Naranjo et al., 2012) under abiotic stresses (cadmium and ethanol respectively). We found that bacterial communities showed more significant responses to melatonin under abiotic stress conditions compared to fungi, along with more distinct separation of communities per treatment (e.g. soil W). Previous soil studies have also found soil bacterial communities to be more responsive to various stress treatments in comparison to fungal communities, with fungi showing greater tolerance to abiotic stressors (Hiroki, 1992, Müller et al., 2001, Marschner et al., 2003, Rajapaksha et al., 2004).

The different responses of soil microbes across the three agricultural soils to melatonin and/or stressors may also have been in part due to differing soil physiochemical characteristics between the soils, as well as differing interactive effects of treatments with various soil characteristics (Zhong and Cai, 2007, Ahn et al., 2012, Zhao et al., 2014, Geisseler et al., 2017). In the current study, some communities were far less impacted by abiotic stress upon the availability of exogenous melatonin. However, as this trend was not observed under cadmium stress, it may be possible that melatonin was utilised by soil bacteria to sustain natural microbial activity by coping with impacts specific to salt stress, such as enhanced osmotic pressure and ion toxicity (Morrissey et al., 2014, Yan et al., 2015).

Melatonin has been consistently shown to reduce cellular levels of ROS in plants exposed to abiotic stress (Tan et al., 2012), by either acting as a highly efficient antioxidant (Reiter et al., 2015, Reiter et al., 2016), or as a signalling molecule, resulting in the upregulation of gene expression, or increased enzyme activities of ROS-scavenging enzymes (Rodriguez et al., 2004, Lee et al., 2014, Manchester et al., 2015, Zhang et al., 2015). As a result, melatonin enhances plant tolerance to abiotic and biotic stresses such as heat, cold, drought and soil contamination as well bacterial and fungal pathogens (Arnao and Hernández-Ruiz, 2013, Arnao and Hernandez-Ruiz, 2014, Zhang et al., 2015, Hardeland, 2016). Melatonin has also been found to increase plant yield by acting as a biostimulator for seed germination and plant growth. As melatonin is safe for human consumption and can be applied to plants in numerous ways such as seed coating, foliar spraying or soil treatment, it may therefore have a major role in future agricultural practices for crop yield protection and improvement (Janas and Posmyk, 2013, Wei et al., 2015, Cui et al., 2017, Zhang et al., 2017).

## Conclusion

In conclusion, this study has demonstrated that exogenous melatonin altered the structures of soil bacterial and, to a lesser extent, fungal assemblages under unstressed and abiotic stressed conditions. No previous reports have examined the effects of melatonin on agricultural soil microbial communities under abiotic stress. Further research is required to profile the microbial taxa responsive to melatonin as well as investigate potential functional associations between melatonin with abiotic stress tolerance in microbes. The main factors causing the differences in natural microbial communities between the different soils also requires further analysis. Additional research is also required to determine if specific soil characteristics influence the responses of microbial communities to melatonin. Moreover, studies may explore potential plant-microbe interactions in soil upon the bioavailability of exogenous melatonin. Future studies involving ameliorating plant stress using melatonin should take into account the potential impact of soil microbiota and the subsequent impact on plant-microbe interactions (beneficial as well as pathogenic). Understanding the role of melatonin in soil microbial community dynamics may provide vital information regarding the viability of melatonin application relating to future agricultural practices.

## Acknowledgements

This work was funded by an International Postgraduate Research Scholarship (IPRS) and Australian Postgraduate Award (APA) provided by La Trobe University, Australia. Soils were sampled with the permissions of Rob Binks in Yaloak estate, Bullan, VIC and Gary Clarke in Melbourne Polytechnic Farm, Yan Yean, VIC, Australia.

## References

Achard-Joris, M., Moreau, J. L., Lucas, M., Baudrimont, M., Mesmer-Dudons, N., Gonzalez, P., Boudou, A. and Bourdineaud, J. P. (2007). Role of metallothioneins in superoxide radical generation during copper redox cycling: Defining the fundamental function of metallothioneins. Biochimie 89, 1474–1488. doi: 10.1016/j.biochi.2007.06.005.

Ahn, J. H., Song, J., Kim, B. Y., Kim, M. S., Joa, J. H. and Weon, H. Y. (2012). Characterization of the bacterial and archaeal communities in rice field soils subjected to long-term fertilization practices. Journal of Microbiology 50, 754–765. doi: 10.1007/s12275-012-2409-6.

Arnao, M. B. and Hernández-Ruiz, J. (2013). Growth conditions determine different melatonin levels in *Lupinus albus* L. J. Pineal Res. 55, 149–155. doi: 10.1111/jpi.12055.

Arnao, M. B. and Hernandez-Ruiz, J. (2014). Melatonin: plant growth regulator and/or biostimulator during stress? Trends Plant Sci. 19, 789–797. doi: 10.1016/j.tplants.2014.07.006.

Arnao, M. B. and Hernández-Ruiz, J. (2015). Functions of melatonin in plants: a review. J. Pineal Res. 59, 133–150. doi: 10.1111/jpi.12253.

Aye, N. S., Sale, P. W. G. and Tang, C. (2016). The impact of long-term liming on soil organic carbon and aggregate stability in low-input acid soils. Biology and Fertility of Soils 52, 697–709. doi: 10.1007/s00374-016-1111-y.

Badri, D. V. and Vivanco, J. M. (2009). Regulation and function of root exudates. Plant, Cell and Environment 32, 666–681. doi: 10.1111/j.1365-3040.2009.01926.x.

Bencherif, K., Boutekrabt, A., Fontaine, J., Laruelle, F., Dalpè, Y. and Anissa, L. H. S. (2015). Impact of soil salinity on arbuscular mycorrhizal fungi biodiversity and microflora biomass associated with *Tamarix articulata* Vahll rhizosphere in arid and semi-arid Algerian areas. Science of the Total Environment 533, 488–494. doi: 10.1016/j.scitotenv.2015.07.007.

Blaalid, R., Kumar, S., Nilsson, R. H., Abarenkov, K., Kirk, P. M. and Kauserud, H. (2013). ITS1 versus ITS2 as DNA metabarcodes for fungi. Molecular Ecology Resources 13, 218–224. doi: 10.1111/1755-0998.12065.

Burns, J. H., Anacker, B. L., Strauss, S. Y. and Burke, D. J. (2015). Soil microbial community variation correlates most strongly with plant species identity, followed by soil chemistry, spatial location and plant genus. AoB PLANTS 7. doi: 10.1093/aobpla/plv030.

Butterly, C. R., Phillips, L. A., Wiltshire, J. L., Franks, A. E., Armstrong, R. D., Chen, D., Mele, P. M. and Tang, C. (2016). Long-term effects of elevated CO_2_ on carbon and nitrogen functional capacity of microbial communities in three contrasting soils. Soil Biology and Biochemistry 97, 157–167. doi: 10.1016/j.soilbio.2016.03.010.

Byeon, Y., Lee, H. Y., Hwang, O. J., Lee, H. J., Lee, K. and Back, K. (2015). Coordinated regulation of melatonin synthesis and degradation genes in rice leaves in response to cadmium treatment. J. Pineal Res. 58, 470–478. doi: 10.1111/jpi.12232.

Cáliz, J., Montserrat, G., Martí, E., Sierra, J., Chung, A. P., Morais, P. V. and Vila, X. (2013). Emerging resistant microbiota from an acidic soil exposed to toxicity of Cr, Cd and Pb is mainly influenced by the bioavailability of these metals. Journal of Soils and Sediments 13, 413–428. doi: 10.1007/s11368-012-0609-7.

Castro, H. F., Classen, A. T., Austin, E. E., Norby, R. J. and Schadt, C. W. (2010). Soil microbial community responses to multiple experimental climate change drivers. Applied and Environmental Microbiology 76, 999–1007. doi: 10.1128/AEM.02874-09.

Chen, C. Q., Fichna, J., Bashashati, M., Li, Y. Y. and Storr, M. (2011). Distribution, function and physiological role of melatonin in the lower gut. World Journal of Gastroenterology 17, 3888–3898. doi: 10.3748/wjg.v17.i34.3888.

Chow, C. E. T., Sachdeva, R., Cram, J. A., Steele, J. A., Needham, D. M., Patel, A., Parada, A. E. and Fuhrman, J. A. (2013). Temporal variability and coherence of euphotic zone bacterial communities over a decade in the Southern California Bight. ISME Journal 7, 2259–2273. doi: 10.1038/ismej.2013.122.

Cui, G., Zhao, X., Liu, S., Sun, F., Zhang, C. and Xi, Y. (2017). Beneficial effects of melatonin in overcoming drought stress in wheat seedlings. Plant Physiol. Biochem. 118, 138–149. doi: 10.1016/j.plaphy.2017.06.014.

Fisher, M. M. and Triplett, E. W. (1999). Automated approach for ribosomal intergenic spacer analysis of microbial diversity and its application to freshwater bacterial communities. Applied and Environmental Microbiology 65, 4630–4636. doi: n/a.

Fuhrman, J. A., Steele, J. A., Hewson, I., Schwalbach, M. S., Brown, M. V., Green, J. L. and Brown, J. H. (2008). A latitudinal diversity gradient in planktonic marine bacteria. Proceedings of the National Academy of Sciences of the United States of America 105, 7774–7778. doi: 10.1073/pnas.0803070105.

Geisseler, D., Linquist, B. A. and Lazicki, P. A. (2017). Effect of fertilization on soil microorganisms in paddy rice systems – A meta-analysis. Soil Biology and Biochemistry 115, 452–460. doi: 10.1016/j.soilbio.2017.09.018.

Gu, Q., Chen, Z., Yu, X., Cui, W., Pan, J., Zhao, G., Xu, S., Wang, R. and Shen, W. (2017). Melatonin confers plant tolerance against cadmium stress via the decrease of cadmium accumulation and reestablishment of microRNA-mediated redox homeostasis. Plant Science 261, 28–37. doi: 10.1016/j.plantsci.2017.05.001.

Hardeland, R. (2015). Melatonin in plants and other phototrophs: advances and gaps concerning the diversity of functions. J. Exp. Bot. 66, 627–646. doi: 10.1093/jxb/eru386.

Hardeland, R. (2016). Melatonin in plants – Diversity of levels and multiplicity of functions. Frontiers in Plant Science 7. doi: 10.3389/fpls.2016.00198.

He, Y., DeSutter, T., Prunty, L., Hopkins, D., Jia, X. and Wysocki, D. A. (2012). Evaluation of 1:5 soil to water extract electrical conductivity methods. Geoderma 185–186, 12–17. doi: 10.1016/j.geoderma.2012.03.022.

Heijboer, A., De Ruiter, P. C., Bodelier, P. L. E. and Kowalchuk, G. A. (2018). Modulation of litter decomposition by the soil microbial food web under influence of land use change. Frontiers in Microbiology 9. doi: 10.3389/fmicb.2018.02860.

Hiroki, M. (1992). Effects of Heavy Metal Contamination on Soil Microbial Population. Soil Science and Plant Nutrition 38, 141–147. doi: 10.1080/00380768.1992.10416961.

Hossain, S. T., Mallick, I. and Mukherjee, S. K. (2012). Cadmium toxicity in Escherichia coli: Cell morphology, Z-ring formation and intracellular oxidative balance. Ecotoxicology and Environmental Safety 86, 54–59. doi: 10.1016/j.ecoenv.2012.09.017.

Janas, K. M. and Posmyk, M. M. (2013). Melatonin, an underestimated natural substance with great potential for agricultural application. Acta Physiol. Plant. 35, 3285–3292. doi: 10.1007/s11738-013-1372-0.

Jiang, C., Cui, Q., Feng, K., Xu, D., Li, C. and Zheng, Q. (2016). Melatonin improves antioxidant capacity and ion homeostasis and enhances salt tolerance in maize seedlings. Acta Physiol. Plant. 38. doi: 10.1007/s11738-016-2101-2.

Jiao, W., Chen, W., Chang, A. C. and Page, A. L. (2012). Environmental risks of trace elements associated with long-term phosphate fertilizers applications: A review. Environmental Pollution 168, 44–53. doi: 10.1016/j.envpol.2012.03.052.

Ke, Q., Kim, H. S., Wang, Z., et al. (2017). Down-regulation of GIGANTEA-like genes increases plant growth and salt stress tolerance in poplar. Plant Biotechnology Journal 15, 331–343. doi: 10.1111/pbi.12628.

Kelly, J. J., Häggblom, M. and Tate Iii, R. L. (1999). Changes in soil microbial communities over time resulting from one time application of zinc: A laboratory microcosm study. Soil Biology and Biochemistry 31, 1455–1465. doi: 10.1016/S0038-0717(99)00059-0.

Khairy, M., El-Safty, S. A. and Shenashen, M. A. (2014). Environmental remediation and monitoring of cadmium. TrAC - Trends in Analytical Chemistry 62, 56–68. doi: 10.1016/j.trac.2014.06.013.

Kirk, J. L., Beaudette, L. A., Hart, M., Moutoglis, P., Klironomos, J. N., Lee, H. and Trevors, J. T. (2004). Methods of studying soil microbial diversity. Journal of Microbiological Methods 58, 169–188. doi: 10.1016/j.mimet.2004.04.006.

Lee, H. Y., Byeon, Y. and Back, K. (2014). Melatonin as a signal molecule triggering defense responses against pathogen attack in *Arabidopsis* and tobacco. J. Pineal Res. 57, 262–268. doi: 10.1111/jpi.12165.

Li, C., Wang, P., Wei, Z., Liang, D., Liu, C., Yin, L., Jia, D., Fu, M. and Ma, F. (2012). The mitigation effects of exogenous melatonin on salinity-induced stress in *Malus hupehensis*. J. Pineal Res. 53, 298–306. doi: 10.1111/j.1600-079X.2012.00999.x.

Li, C., Zhao, Q., Gao, T., Wang, H., Zhang, Z., Liang, B., Wei, Z., Liu, C. and Ma, F. (2018). The mitigation effects of exogenous melatonin on replant disease in apple. J. Pineal Res. 65. doi: 10.1111/jpi.12523.

Li, M. Q., Hasan, M. K., Li, C. X., et al. (2016). Melatonin mediates selenium-induced tolerance to cadmium stress in tomato plants. J. Pineal Res. 291–302. doi: 10.1111/jpi.12346.

Liang, C., Zheng, G., Li, W., et al. (2015). Melatonin delays leaf senescence and enhances salt stress tolerance in rice. J. Pineal Res. 59, 91–101. doi: 10.1111/jpi.12243.

Liu, T., Zhao, F., Liu, Z., Zuo, Y., Hou, J. and Wang, Y. (2016). Identification of melatonin in *Trichoderma* spp. and detection of melatonin content under controlled-stress growth conditions from *T. asperellum*. Journal of Basic Microbiology 56, 838–843. doi: 10.1002/jobm.201500223.

Lucchelli, A., Santagostino-Barbone, M. G. and Tonini, M. (1997). Investigation into the contractile response of melatonin in the guinea-pig isolated proximal colon: The role of 5-HT4 and melatonin receptors. British Journal of Pharmacology 121, 1775–1781. doi: 10.1038/sj.bjp.0701287.

Manchester, L. C., Coto-Montes, A., Boga, J. A., Andersen, L. P. H., Zhou, Z., Galano, A., Vriend, J., Tan, D. X. and Reiter, R. J. (2015). Melatonin: An ancient molecule that makes oxygen metabolically tolerable. J. Pineal Res. 59, 403–419. doi: 10.1111/jpi.12267.

Marschner, P., Kandeler, E. and Marschner, B. (2003). Structure and function of the soil microbial community in a long-term fertilizer experiment. Soil Biology and Biochemistry 35, 453–461. doi: 10.1016/S0038-0717(02)00297-3.

Mau, R. L., Dijkstra, P., Schwartz, E., Koch, B. J. and Hungate, B. A. (2019). Addressing strengths and weaknesses of a multi-ecosystem climate change experiment. Applied Soil Ecology 133, 191–192. doi: 10.1016/j.apsoil.2018.07.003.

Morrissey, E. M., Gillespie, J. L., Morina, J. C. and Franklin, R. B. (2014). Salinity affects microbial activity and soil organic matter content in tidal wetlands. Global Change Biology 20, 1351–1362. doi: 10.1111/gcb.12431.

Müller, A. K., Westergaard, K., Christensen, S. and Sørensen, S. J. (2001). The effect of long-term mercury pollution on the soil microbial community. FEMS Microbiology Ecology 36, 11–19. doi: 10.1016/S0168-6496(01)00112-X.

Ondreičková, K., Žofajová, A., Piliarová, M., Gubiš, J. and Hudcovicová, M. (2016). Monitoring of rhizosphere bacterial communities in soil with sewage sludge addition using two molecular fingerprinting methods: Do these methods give similar results? Agriculture 62, 52–61. doi: 10.1515/agri-2016-0006.

Öztürk, A. I., Yilmaz, O., Kirbağ, S. and Arslan, M. (2000). Antimicrobial and biological effects of ipemphos and amphos on bacterial and yeast strains. Cell Biochemistry and Function 18, 117–126. doi: 10.1002/(SICI)1099-0844(200006)18:2<117::AID-CBF863>3.0.CO;2-1.

Paulose, J. K., Wright, J. M., Patel, A. G. and Cassone, V. M. (2016). Human gut bacteria are sensitive to melatonin and express endogenous circadian rhythmicity. PLoS One 11. doi: 10.1371/journal.pone.0146643.

Rajapaksha, R. M. C. P., Tobor-Kapłon, M. A. and Bååth, E. (2004). Metal toxicity affects fungal and bacterial activities in soil differently. Applied and Environmental Microbiology 70, 2966–2973. doi: 10.1128/AEM.70.5.2966-2973.2004.

Ramette, A. (2009). Quantitative community fingerprinting methods for estimating the abundance of operational taxonomic units in natural microbial communities. Applied and Environmental Microbiology 75, 2495–2505. doi: 10.1128/AEM.02409-08.

Rath, K. M., Maheshwari, A., Bengtson, P. and Rousk, J. (2016). Comparative toxicities of salts on microbial processes in soil. Applied and Environmental Microbiology 82, 2012–2020. doi: 10.1128/AEM.04052-15.

Reese, A. T., Lulow, K., David, L. A. and Wright, J. P. (2018). Plant community and soil conditions individually affect soil microbial community assembly in experimental mesocosms. Ecology and Evolution 8, 1196–1205. doi: 10.1002/ece3.3734.

Reiter, R. J., Tan, D. X., Zhou, Z., Cruz, M. H. C., Fuentes-Broto, L. and Galano, A. (2015). Phytomelatonin: Assisting plants to survive and thrive. Molecules 20, 7396–7437. doi: 10.3390/molecules20047396.

Reiter, R. J., Mayo, J. C., Tan, D. X., Sainz, R. M., Alatorre-Jimenez, M. and Qin, L. (2016). Melatonin as an antioxidant: under promises but over delivers. J. Pineal Res. 61, 253–278. doi: 10.1111/jpi.12360.

Rincon-Florez, V. A., Carvalhais, L. C. and Schenk, P. M. (2013). Culture-independent molecular tools for soil and rhizosphere microbiology. Diversity 5, 581–612. doi: 10.3390/d5030581.

Roberts, T. L. (2014). Cadmium and phosphorous fertilizers: The issues and the science. Procedia Engineering 83, 52–59. doi: 10.1016/j.proeng.2014.09.012

Rodriguez-Naranjo, M. I., Torija, M. J., Mas, A., Cantos-Villar, E. and Garcia-Parrilla, M. D. C. (2012). Production of melatonin by *Saccharomyces* strains under growth and fermentation conditions. J. Pineal Res. 53, 219–224. doi: 10.1111/j.1600-079X.2012.00990.x.

Rodriguez, C., Mayo, J. C., Sainz, R. M., Antolín, I., Herrera, F., Martín, V. and Reiter, R. J. (2004). Regulation of antioxidant enzymes: A significant role for melatonin. J. Pineal Res. 36, 1–9. doi: 10.1046/j.1600-079X.2003.00092.x.

Schultz, C. L., Edrington, T. S., Callaway, T. R., Schroeder, S. B., Hallford, D. M., Genovese, K. J., Anderson, R. C. and Nisbet, D. J. (2006). The influence of melatonin on growth of *E. coli* O157:H7 in pure culture and exogenous melatonin on faecal shedding of *E. coli* O157:H7 in experimentally infected wethers. Letters in Applied Microbiology 43, 105–110. doi: 10.1111/j.1472-765X.2006.01909.x.

Sørensen, J., Nicolaisen, M. H., Ron, E. and Simonet, P. (2009). Molecular tools in rhizosphere microbiology-from single-cell to whole-community analysis. Plant and Soil 321, 483–512. doi: 10.1007/s11104-009-9946-8.

Tan, D. X., Hardeland, R., Manchester, L. C., Korkmaz, A., Ma, S., Rosales-Corral, S. and Reiter, R. J. (2012). Functional roles of melatonin in plants, and perspectives in nutritional and agricultural science. J. Exp. Bot. 63, 577–597. doi: 10.1093/jxb/err256.

Tan, D. X., Zheng, X. D., Kong, J., Manchester, L. C., Hardeland, R., Kim, S. J., Xu, X. Y. and Reiter, R. J. (2014). Fundamental issues related to the origin of melatonin and melatonin isomers during evolution: relation to their biological functions. Int. J. Mol. Sci. 15, 15858–15890. doi: 10.3390/ijms150915858.

Tanaka, T., Nishio, K., Usuki, Y. and Fujita, K. i. (2006). Involvement of oxidative stress induction in Na^+^ toxicity and its relation to the inhibition of a Ca^2+^-dependent but calcineurin-independent mechanism in *Saccharomyces cerevisiae*. Journal of Bioscience and Bioengineering 101, 77–79. doi: 10.1263/jbb.101.77.

Tekbas, O. F., Ogur, R., Korkmaz, A., Kilic, A. and Reiter, R. J. (2008). Melatonin as an antibiotic: New insights into the actions of this ubiquitous molecule. J. Pineal Res. 44, 222–226. doi: 10.1111/j.1600-079X.2007.00516.x.

Van Wijngaarden, R. P. A., Van Den Brink, P. J., Oude Voshaar, J. H. and Leeuwangh, P. (1995). Ordination techniques for analysing response of biological communities to toxic stress in experimental ecosystems. Ecotoxicology 4, 61–77. doi: 10.1007/BF00350650.

Wahid, F., Sharif, M., Steinkellner, S., Khan, M. A., Marwat, K. B. and Khan, S. A. (2016). Inoculation of arbuscular mycorrhizal fungi and phosphate solubilizing bacteria in the presence of rock phosphate improves phosphorus uptake and growth of maize. Pakistan Journal of Botany 48, 739–747. doi.

Wall, D. H. and Virginia, R. A. (1999). Controls on soil biodiversity: Insights from extreme environments. Applied Soil Ecology 13, 137–150. doi: 10.1016/S0929-1393(99)00029-3.

Wang, H. X., Liu, F. and Ng, T. B. (2001). Examination of pineal indoles and 6-methoxy-2-benzoxazolinone for antioxidant and antimicrobial effects. Comparative Biochemistry and Physiology - C Toxicology and Pharmacology 130, 379–388. doi: 10.1016/S1532-0456(01)00264-2.

Weeda, S., Zhang, N., Zhao, X. L., Ndip, G., Guo, Y. D., Buck, G. A., Fu, C. G. and Ren, S. X. (2014). *Arabidopsis* transcriptome analysis reveals key roles of melatonin in plant defense systems. PLoS One 9, 18. doi: 10.1371/journal.pone.0093462.

Wei, W., Li, Q. T., Chu, Y. N., et al. (2015). Melatonin enhances plant growth and abiotic stress tolerance in soybean plants. J. Exp. Bot. 66, 695–707. doi: 10.1093/jxb/eru392.

Wood, J. L., Tang, C. and Franks, A. E. (2016). Microbial associated plant growth and heavy metal accumulation to improve phytoextraction of contaminated soils. Soil Biology and Biochemistry 103, 131–137. doi: 10.1016/j.soilbio.2016.08.021.

Wood, J. L., Zhang, C., Mathews, E. R., Tang, C. and Franks, A. E. (2016). Microbial community dynamics in the rhizosphere of a cadmium hyper-accumulator. Scientific Reports 6, 1–10. doi: 10.1038/srep36067.

Yan, N., Marschner, P., Cao, W., Zuo, C. and Qin, W. (2015). Influence of salinity and water content on soil microorganisms. International Soil and Water Conservation Research 3, 316–323. doi: 10.1016/j.iswcr.2015.11.003.

Yao, H., He, Z., Wilson, M. J. and Campbell, C. D. (2000). Microbial biomass and community structure in a sequence of soils with increasing fertility and changing land use. Microbial Ecology 40, 223–237. doi: n/a

Zancarini, A., Mougel, C., Voisin, A. S., Prudent, M., Salon, C. and Munier-Jolain, N. (2012). Soil nitrogen availability and plant genotype modify the nutrition strategies of *M. truncatula* and the associated rhizosphere microbial communities. PLoS One 7. doi: 10.1371/journal.pone.0047096.

Zhang, N., Sun, Q., Zhang, H., Cao, Y., Weeda, S., Ren, S. and Guo, Y. D. (2015). Roles of melatonin in abiotic stress resistance in plants. J. Exp. Bot. 66, 647–656. doi: 10.1093/jxb/eru336.

Zhang, Q., Jin, H., Zhou, H., Cai, M., Li, Y., Zhang, G. and Di, H. (2019). Variation of soil anaerobic microorganisms connected with anammox processes by 13C-phospholipid fatty acid analysis among long-term fertilization regimes in a crop rotation system. Applied Soil Ecology 133, 34–43. doi: 10.1016/j.apsoil.2018.09.005.

Zhang, Y. P., Yang, S. J. and Chen, Y. Y. (2017). Effects of melatonin on photosynthetic performance and antioxidants in melon during cold and recovery. Biologia Plantarum 61, 571–578. doi: 10.1007/s10535-017-0717-8.

Zhao, J., Ni, T., Li, Y., Xiong, W., Ran, W., Shen, B., Shen, Q. and Zhang, R. (2014). Responses of bacterial communities in arable soils in a rice-wheat cropping system to different fertilizer regimes and sampling times. PLoS One 9. doi: 10.1371/journal.pone.0085301.

Zhong, W. H. and Cai, Z. C. (2007). Long-term effects of inorganic fertilizers on microbial biomass and community functional diversity in a paddy soil derived from quaternary red clay. Applied Soil Ecology 36, 84–91. doi: 10.1016/j.apsoil.2006.12.001.

