## Supporting Information for "Bacterial and fungal communities are differentially modified by melatonin in agricultural soils under abiotic stress"

### SUPPLEMENTARY INFORMATION

**Supplementary Table 1**: PERMANOVA analyses of a) bacterial and b) fungal community responses to treatments with high and low stresses based on ARISA data (Bray-Curtis dissimilarity distances).

**a)**

| | Soil P | | | |
|---|---|---|---|---|
| | **Cadmium** | | **Salt** | |
| | **t-statistic** | **P-value** | **t-statistic** | **P-value** |
| H-stress vs L-stress | 1.0771 | 0.2882 | 2.2813 | **0.0003*** ** |
| H-stress vs Control | 1.4669 | 0.073 | 1.3298 | **0.0431*** |
| L-stress vs Control | 1.1248 | 0.2325 | 1.6868 | **0.0089**** |

| | Soil C | | | |
|---|---|---|---|---|
| | **Cadmium** | | **Salt** | |
| | **t-statistic** | **P-value** | **t-statistic** | **P-value** |
| H-stress vs L-stress | 2.7886 | **0.0001***** | 5.0877 | **0.0001***** |
| H-stress vs Control | 4.0829 | **0.0001***** | 5.861 | **0.0001***** |
| L-stress vs Control | 1.933 | **0.0035**** | 1.684 | **0.0158*** |

| | Soil W | | | |
|---|---|---|---|---|
| | **Cadmium** | | **Salt** | |
| | **t-statistic** | **P-value** | **t-statistic** | **P-value** |
| H-stress vs L-stress | 2.3933 | **0.0001***** | 3.1672 | **0.0001***** |
| H-stress vs Control | 3.9566 | **0.0001***** | 3.9779 | **0.0001***** |
| L-stress vs Control | 2.2386 | **0.0001***** | 2.3602 | **0.0001***** |

**b)**

| Soil P | Cadmium | | Salt | |
|---|---|---|---|---|
| | **t-statistic** | **P-value** | **t-statistic** | **P-value** |
| H-stress vs L-stress | 1.026 | 0.3837 | 1.7222 | **0.0031**** |
| H-stress vs Control | 1.8378 | **0.003**** | 1.3462 | **0.036*** |
| L-stress vs Control | 2.0151 | **0.0008***** | 1.059 | 0.3396 |

| Soil C | Cadmium | | Salt | |
|---|---|---|---|---|
| | **t-statistic** | **P-value** | **t-statistic** | **P-value** |
| H-stress vs L-stress | 1.7138 | **0.0076**** | 1.4761 | **0.0255*** |
| H-stress vs Control | 1.5488 | **0.0145*** | 2.8112 | **0.0002***** |
| L-stress vs Control | 1.1351 | 0.23 | 1.8004 | **0.0009***** |

| Soil W | Cadmium | | Salt | |
|---|---|---|---|---|
| | **t-statistic** | **P-value** | **t-statistic** | **P-value** |
| H-stress vs L-stress | 1.6613 | **0.0255*** | 2.0967 | **0.0002***** |
| H-stress vs Control | 1.5842 | **0.0103*** | 3.2649 | **0.0001***** |
| L-stress vs Control | 0.99408 | 0.4411 | 1.6346 | **0.0131*** |

Treatments: H-stress = High stressor; L-stress = Low stressor; Control = MQ water replacing stressor. All treatments and controls for the same soil were composed of a standardised amount of dilute ethanol. n=4 replicates per treatment. Significance of PERMANOVA (highlighted in bold): *: 0.01 < p-value ≤ 0.05; **: 0.001 < p-value ≤ 0.01; ***: p-value ≤ 0.001.

**Supplementary Data 1**

**Effects of treatments on α diversity (OTU richness)**

Community fingerprinting with ARISA allowed for a statistical estimation of the α diversity based on the different Operational Taxonomic Units (OTUs) observed. The basal levels of α diversity determined in the dry, untreated samples from P, C and W indicated a mean of 29.7, 30 and 36 bacterial OTUs, and 16.6, 16.4 and 14.8 fungal OTUs, respectively. Concurrently, across all three control soil samples, the number of OTUs observed for bacterial communities (23-39) were considerably greater than the OTU numbers for fungi (8-20). Such findings are in line with other studies where community fingerprinting with ARISA detected substantially fewer fungal OTUs compared to bacterial (Ranjard et al., 2001, Hansgate et al., 2005).

**Melatonin alters microbial α diversity**

In melatonin-treated soils the mean bacterial OTU richness (α diversity) was 29.9, 34.3 and 34.2 in soils P, C and W (ranging from 21 to 40 OTUs; Supplementary Table 2a) and 16.1, 14.4 and 13 (ranging from 9 to 19 OTUs; Supplementary Table 2b) for fungi respectively. The effect of melatonin treatment at high (H) and low (L) concentrations on OTU numbers varied for bacterial communities across the three soils compared to control samples (Supplementary Table 2a). Bacterial α diversities shifted ($p < 0.05$) in response to high melatonin treatment within all three soils, whereas low melatonin treatment impacted bacterial community only in soil C. At sampling timepoints T0 and T1, bacterial OTU richness's were significantly ($p < 0.05$) decreased compared to respective control samples upon treatment with high melatonin for soil W. High melatonin also resulted in significant decrease in bacterial OTU numbers in soil P relative to controls at sampling timepoint T0, but not T1. Interestingly bacterial OTU richness in high and low melatonin-treated soils for soil C significantly ($p < 0.01$) increased at T1, but no significant changes were observed at T0. In contrast to the varying responses of bacterial assemblages, OTU richness of fungal communities across all three soils was not impacted ($p > 0.05$) by melatonin (Supplementary Table 2b).

The number of bacterial OTUs common to control and melatonin-only treatments varied considerably for each soil between the sampling timepoints. For example, at timepoint T0, 73.1% (49/67), 73.7% (45/61) and 73.2% (52/71) of bacterial OTUs were common to control and melatonin-only treatments in soils P, C and W respectively. However, these numbers reduced at timepoint T1 to 56.4% (44/78), 43.6% (34/78) and 55.3% (42/76) respectively.

Fungal OTUs common to control and melatonin-only treatments in soils varied less compared to bacterial assemblages. At sampling timepoint T0, 75.8% (22/29), 71.4% (20/28) and 76.2% (16/21) fungal OTUs were common to control and melatonin-only treatments in soils P, C and W respectively. At sampling timepoint T0, the common OTUs reduced to 63.6% (21/33) for soil P, whereas OTU richness increased slightly to 74.1% (20/27) and 80.8% (21/26) for soils C and W respectively.

**Cadmium and salt show limited impacts on microbial α diversity**

OTU richness (α diversity) was not impacted by stressor level (high [H] or low [L]) in cadmium or salt experiment (PERMANOVA, $p > 0.05$) for bacterial communities in soils P and W (Supplementary Table 2a). For soil C, low salt treatment significantly increased in OTU numbers ($p < 0.001$) relative to control. OTU numbers for fungal communities across all soils were not impacted ($p > 0.05$) by either cadmium or salt stresses (Supplementary Table 2b).

The number of bacterial OTUs common to cadmium-only treatments and the control samples for each soil were relatively consistent within soils P, C and W at 63.0% (46/73), 60.3% (41/68) and 59.5% (44/74) respectively. In contrast, bacterial OTU numbers common to salt-only and control treatments showed greater variation at 67.6% (48/71), 45.3% (34/75) and 60.8% (45/74) for soils P, C and W respectively. Fungal OTUs common to cadmium-only treatments and the control samples were varied for soils P, C and W at 73.3% (22/30), 68.0% (17/25) and 84.2% (16/19) respectively. In comparison, fungal OTUs for salt-only and control treatments showed greater consistency with 75.8% (22/29), 71.4% (20/28) and 75.8% (22/29) common across soils P, C and W respectively.

**Supplementary Table 2**: Mean OTU richness (α diversity) (± SE) within a) bacterial and b) fungal communities for various melatonin- or stressor-only treatments (n=4 replicates). MT = Melatonin (High concentration = 4 mg/kg soil; Low concentration = 0.2 mg/kg soil). Stressors: T0 = cadmium (High concentration = 280 mg/kg soil; Low concentration = 100 mg/kg soil); T1 = salt (High concentration = 7 g/kg soil; Low concentration = 4 g/kg soil). Melatonin-treated soils contained no abiotic stressor. Control = dilute ethanol solution (i.e. no melatonin, no stressor).

**a)**

| | **Soil P** | | **Soil C** | | **Soil W** | |
|---|---|---|---|---|---|---|
| | **T0** | **T1** | **T0** | **T1** | **T0** | **T1** |
| High MT | **24.0 ± 1.5** * | 33.5 ± 2.3 | 33.8 ± 1.7 | **36.3 ± 0.9** * | **28.8 ± 1.0** * | **34.3 ± 0.3** * |
| Low MT | 28.8 ± 1.1 | 33.5 ± 2.8 | 30.8 ± 2.8 | **36.5 ± 0.9** * | 37.5 ± 0.9 | 36.3 ± 1.0 |
| High stress | 31.0 ± 0.4 | 33.8 ± 0.8 | 28.3 ± 1.9 | 22.5 ± 0.6 | 34.0 ± 0.4 | 37.5 ± 0.5 |
| Low stress | 31.3 ± 0.8 | 32.3 ± 0.8 | 28.8 ± 1.1 | **36.5 ± 1.3** * | 34.3 ± 0.5 | 37.5 ± 0.6 |
| Control | 31.5 ± 1.0 | 31.8 ± 0.8 | 33.8 ± 1.7 | 26.0 ± 1.1 | 37.0 ± 0.4 | 38.3 ± 0.8 |

**b)**

| | **Soil P** | | **Soil C** | | **Soil W** | |
|---|---|---|---|---|---|---|
| | **T0** | **T1** | **T0** | **T1** | **T0** | **T1** |
| High MT | 17.3 ± 1.0 | 14.5 ± 1.0 | 16.8 ± 0.8 | 15.0 ± 0.7 | 13.3 ± 0.9 | 11.0 ± 1.1 |
| Low MT | 16.8 ± 0.5 | 15.8 ± 1.5 | 12.8 ± 1.3 | 13.3 ± 0.9 | 16.3 ± 0.6 | 11.8 ± 0.5 |
| High stress | 15.8 ± 1.7 | 17.3 ± 0.5 | 16.0 ± 1.4 | 13.5 ± 0.5 | 11.8 ± 0.5 | 12.3 ± 0.5 |
| Low stress | 18.0 ± 1.1 | 15.8 ± 0.6 | 15.0 ± 1.2 | 14.3 ± 0.5 | 14.3 ± 1.7 | 11.5 ± 0.3 |
| Control | 17.8 ± 1.1 | 16.0 ± 1.2 | 16.0 ± 0.9 | 13.0 ± 0.6 | 15.0 ± 0.9 | 10.8 ± 1.1 |

All treatments and controls in the same soil were composed of a standardised amount of dilute ethanol. OTU numbers in bold were determined as being significantly different to control samples using one-way ANOVA. * indicates significance level of $p < 0.01$ (highlighted in bold and shaded).

**Supplementary Table 3:** The largest increases of relative abundance (%) of individual OTUs from melatonin-only treated samples compared with controls as determined by ANOSIM.

**Bacteria**

| Sampling Timepoint | MT treatment | Soil | OTU | MT abundance (%) | Control Abundance (%) | Difference (%) |
|---|---|---|---|---|---|---|
| T1 | High | W | 741.4 | 7.09 | 0 | 7.1 |
| T1 | Low | W | 741.4 | 6.55 | 0 | 6.6 |
| T0 | High | P | 447 | 16.98 | 10.44 | 6.5 |
| T0 | High | W | 447 | 10.54 | 4.13 | 6.4 |
| T1 | Low | C | 741.4 | 8.34 | 2.21 | 6.1 |
| T1 | High | W | 501.4 | 8.22 | 2.2 | 6.0 |
| T1 | High | C | 741.4 | 7.31 | 2.21 | 5.1 |
| T0 | High | W | 183 | 9.12 | 4.36 | 4.8 |
| T1 | High | C | 447.4 | 8.64 | 4.27 | 4.4 |
| T1 | Low | C | 384.4 | 4.27 | 0 | 4.3 |
| T1 | Low | C | 501.4 | 6.92 | 2.81 | 4.1 |
| T1 | High | W | 183.4 | 5.58 | 1.9 | 3.7 |
| T1 | Low | W | 501.4 | 5.75 | 2.2 | 3.6 |

**Fungi**

| Sampling Timepoint | MT treatment | Soil | OTU | MT abundance (%) | Control Abundance (%) | Difference (%) |
|---|---|---|---|---|---|---|
| T0 | High | W | 607.5 | 17.33 | 5.8 | 11.5 |
| T1 | High | W | 612.5 | 7.63 | 0.92 | 6.7 |
| T0 | Low | C | 599.5 | 7.24 | 1.28 | 6.0 |
| T0 | High | W | 675.5 | 9.55 | 4.23 | 5.3 |
| T0 | Low | P | 567.5 | 11.79 | 7.21 | 4.6 |
| T0 | High | C | 575.5 | 45.02 | 40.96 | 4.1 |
| T0 | High | P | 559.5 | 7.89 | 3.95 | 3.9 |
| T0 | Low | C | 591.5 | 11.69 | 8.11 | 3.6 |
| T1 | High | W | 596.5 | 21.24 | 17.84 | 3.4 |
| T0 | Low | W | 607.5 | 8.96 | 5.8 | 3.2 |
| T1 | High | P | 676.5 | 5.19 | 2.13 | 3.1 |
| T1 | Low | C | 576.5 | 37.42 | 34.42 | 3.0 |
| T1 | Low | P | 708.5 | 2.74 | 0 | 2.7 |

MT – Melatonin; OTU: Operational Taxonomic Unit; Soil P: Pasture; Soil C: Canola; Soil W: Wheat [+fire blazing]; High MT = 4 mg $kg^{-1}$ soil; Low MT = 0.2 mg $kg^{-1}$ soil.

**Supplementary Table 4:** The largest decreases of relative abundance (%) of individual OTUs from melatonin-only treated samples compared with controls as determined by ANOSIM.

**Bacteria**

| Sampling Timepoint | MT treatment | Soil | OTU | MT abundance (%) | Control Abundance (%) | Difference (%) |
|---|---|---|---|---|---|---|
| T1 | High | C | 849.4 | 2.42 | 19.34 | -16.9 |
| T1 | Low | C | 756.4 | 2.69 | 18.89 | -16.2 |
| T1 | High | C | 756.4 | 2.79 | 18.89 | -16.1 |
| T1 | Low | C | 849.4 | 3.3 | 19.34 | -16.0 |
| T1 | High | W | 849.4 | 0 | 12.48 | -12.5 |
| T1 | Low | W | 849.4 | 1.28 | 12.48 | -11.2 |
| T1 | Low | W | 756.4 | 0 | 6.9 | -6.9 |
| T1 | High | W | 756.4 | 0.28 | 6.9 | -6.6 |
| T0 | High | W | 339 | 3.06 | 8.56 | -5.5 |
| T1 | Low | P | 156.4 | 2.33 | 6.43 | -4.1 |
| T0 | Low | C | 738 | 4.61 | 8.36 | -3.8 |
| T1 | Low | P | 180.4 | 2.27 | 5.86 | -3.6 |
| T1 | High | P | 180.4 | 2.74 | 5.86 | -3.1 |

**Fungi**

| Sampling Timepoint | MT treatment | Soil | OTU | MT abundance (%) | Control Abundance (%) | Difference (%) |
|---|---|---|---|---|---|---|
| T0 | Low | C | 575.5 | 30.73 | 40.96 | -10.2 |
| T1 | High | W | 616.5 | 0 | 9.6 | -9.6 |
| T0 | High | W | 591.5 | 15.82 | 23.59 | -7.8 |
| T0 | Low | W | 591.5 | 17.63 | 23.59 | -6.0 |
| T0 | High | W | 755.5 | 0.38 | 6.29 | -5.9 |
| T0 | High | W | 575.5 | 27.67 | 33.45 | -5.8 |
| T0 | Low | W | 755.5 | 1.8 | 6.29 | -4.5 |
| T1 | High | P | 576.5 | 31.3 | 35.6 | -4.3 |
| T1 | High | P | 876.5 | 0 | 4.24 | -4.2 |
| T1 | Low | P | 876.5 | 0 | 4.24 | -4.2 |
| T0 | High | C | 607.5 | 6.98 | 11.13 | -4.2 |
| T0 | High | P | 591.5 | 12.71 | 16.72 | -4.0 |
| T1 | Low | C | 584.5 | 1.05 | 4.72 | -3.7 |

MT – melatonin; OTU: Operational Taxonomic Unit; Soil P: Pasture; Soil C: Canola; Soil W: Wheat [+fire blazing]; High MT = 4 mg $kg^{-1}$ soil; Low MT = 0.2 mg $kg^{-1}$ soil.

**Supplementary Table 5**: PERMANOVA analyses of a) bacterial and b) fungal community responses to all treatments with high and low melatonin (i.e. no stressors included) at sampling timepoints T0 and T1, based on ARISA data (Bray-Curtis dissimilarity distances).

**a)**

| | Soil P | | | |
|---|---|---|---|---|
| | T0 | | T1 | |
| | **t-statistic** | **P-value** | **t-statistic** | **P-value** |
| H-MT vs L-MT | 0.98661 | 0.4074 | 1.5173 | **0.0103*** |
| H-MT vs Control | 1.3386 | 0.0928 | 1.7634 | **0.0008***** |
| L-MT vs Control | 1.2234 | 0.1587 | 1.3574 | 0.0665 |

| | Soil C | | | |
|---|---|---|---|---|
| | T0 | | T1 | |
| | **t-statistic** | **P-value** | **t-statistic** | **P-value** |
| H-MT vs L-MT | 1.6225 | **0.0263*** | 1.1227 | 0.2484 |
| H-MT vs Control | 0.8002 | 0.7636 | 1.8497 | **0.0101*** |
| L-MT vs Control | 1.7851 | **0.0135*** | 2.2132 | **0.0029**** |

| | Soil W | | | |
|---|---|---|---|---|
| | T0 | | T1 | |
| | **t-statistic** | **P-value** | **t-statistic** | **P-value** |
| H-MT vs L-MT | 3.5945 | **0.0001***** | 2.8317 | **0.0001***** |
| H-MT vs Control | 4.3751 | **0.0001***** | 3.7099 | **0.0001***** |
| L-MT vs Control | 1.3464 | **0.0261*** | 1.6455 | **0.0006***** |

**b)**

| | Soil P | | | |
|---|---|---|---|---|
| | T0 | | T1 | |
| | **t-statistic** | **P-value** | **t-statistic** | **P-value** |
| H-MT vs L-MT | 1.2699 | 0.1042 | 1.0662 | 0.3265 |
| H-MT vs Control | 0.92172 | 0.5693 | 1.4434 | **0.0194*** |
| L-MT vs Control | 1.0088 | 0.4313 | 1.0374 | 0.3796 |

| | Soil C | | | |
|---|---|---|---|---|
| | T0 | | T1 | |
| | **t-statistic** | **P-value** | **t-statistic** | **P-value** |
| H-MT vs L-MT | 0.90452 | 0.6202 | 0.71054 | 0.9021 |
| H-MT vs Control | 1.1021 | 0.2829 | 1.1387 | 0.2269 |
| L-MT vs Control | 1.063 | 0.3425 | 1.304 | 0.0727 |

| | Soil W | | | |
|---|---|---|---|---|
| | T0 | | T1 | |
| | **t-statistic** | **P-value** | **t-statistic** | **P-value** |
| H-MT vs L-MT | 1.87 | **0.0068**** | 1.7212 | **0.0041**** |
| H-MT vs Control | 2.7675 | **0.0005***** | 1.7703 | **0.0005***** |
| L-MT vs Control | 1.6514 | **0.0248*** | 0.73838 | 0.8333 |

H-MT: High melatonin; L-MT: Low melatonin. Control treatments consisted of dilute ethanol replacing melatonin. All treatments and controls for the same soil were composed of a standardised amount of dilute ethanol. n=4 replicates per treatment. Significance of PERMANOVA (highlighted in bold): *: 0.01 < p-value ≤ 0.05; **: 0.001 < p-value ≤ 0.01; ***: p-value ≤ 0.001.

**Supplementary Table 6**: Differences (a < 0.05) between a) bacterial and b) fungal communities treated with melatonin under various stressor conditions as determined by PERMANOVA using Monte Carlo simulation [P-(MC)]. For control treatments, melatonin was replaced with dilute ethanol.

**a)**

| | Cadmium stress | | | | | | Salt stress | | | | | |
|---|---|---|---|---|---|---|---|---|---|---|---|---|
| | High stressor | | Low stressor | | No stressor | | High stressor | | Low stressor | | No stressor | |
| | t-statistic | P-(MC) | t-statistic | P-(MC) | t-statistic | P-(MC) | t-statistic | P-(MC) | t-statistic | P-(MC) | t-statistic | P-(MC) |
| **Soil P** | | | | | | | | | | | | |
| H-MT vs L-MT | 1.8896 | 0.0381* | 1.7726 | 0.0426* | 0.79519 | 0.5926 | 1.2743 | 0.1795 | 1.0879 | 0.3402 | 1.228 | 0.23 |
| H-MT vs Control | 2.0701 | 0.0174* | 1.0604 | 0.3504 | 1.044 | 0.3492 | 1.8399 | 0.0221* | 1.6486 | 0.0723 | 0.9332 | 0.4988 |
| L-MT vs Control | 1.4365 | 0.1102 | 1.3795 | 0.135 | 1.0178 | 0.3787 | 1.3481 | 0.1393 | 1.2687 | 0.2054 | 1.3996 | 0.1239 |
| **Soil C** | | | | | | | | | | | | |
| H-MT vs L-MT | 0.67148 | 0.7288 | 1.2021 | 0.241 | 1.5997 | 0.0869 | 0.9219 | 0.4359 | 1.8272 | 0.0367* | 1.1973 | 0.2436 |
| H-MT vs Control | 1.3559 | 0.1542 | 0.85514 | 0.5686 | 0.6089 | 0.8136 | 0.92863 | 0.4766 | 3.2053 | 0.0021** | 3.9213 | 0.0006*** |
| L-MT vs Control | 1.5054 | 0.0911 | 0.69004 | 0.7426 | 1.6304 | 0.0763 | 1.2148 | 0.2422 | 2.527 | 0.0072** | 4.1967 | 0.0006*** |
| **Soil W** | | | | | | | | | | | | |
| H-MT vs L-MT | 1.4721 | 0.1032 | 2.502 | 0.0056** | 3.2428 | 0.0017** | 1.3523 | 0.1416 | 2.7151 | 0.0039** | 2.2224 | 0.0106* |
| H-MT vs Control | 2.7945 | 0.0041** | 2.4253 | 0.0071** | 3.0177 | 0.0031** | 2.2808 | 0.0068** | 4.4083 | 0.0004*** | 4.3159 | 0.0005*** |
| L-MT vs Control | 1.9951 | 0.0123* | 1.1457 | 0.2718 | 0.74773 | 0.7544 | 2.4454 | 0.0052** | 3.6857 | 0.0013** | 3.1744 | 0.0031** |

**b)**

| | Cadmium stress | | | | | | Salt stress | | | | | |
|---|---|---|---|---|---|---|---|---|---|---|---|---|
| | High stressor | | Low stressor | | No stressor | | High stressor | | Low stressor | | No stressor | |
| | **t-statistic** | **P-(MC)** | **t-statistic** | **P-(MC)** | **t-statistic** | **P-(MC)** | **t-statistic** | **P-(MC)** | **t-statistic** | **P-(MC)** | **t-statistic** | **P-(MC)** |
| **Soil P** | | | | | | | | | | | | |
| H-MT vs L-MT | 1.1042 | 0.3099 | 1.3741 | 0.1364 | 1.1155 | 0.2973 | 1.2556 | 0.1849 | 0.83979 | 0.616 | 1.2733 | 0.1992 |
| H-MT vs Control | 1.2813 | 0.2003 | 0.764 | 0.6662 | 1.0154 | 0.4022 | 1.4583 | 0.0879 | 0.91786 | 0.5061 | 1.2197 | 0.2311 |
| L-MT vs Control | 0.99918 | 0.4228 | 1.1821 | 0.2647 | 0.64845 | 0.8226 | 1.2441 | 0.1989 | 0.97139 | 0.4403 | 1.1837 | 0.2552 |
| **Soil C** | | | | | | | | | | | | |
| H-MT vs L-MT | 0.60558 | 0.8292 | 0.74193 | 0.7091 | 1.445 | 0.1233 | 0.90493 | 0.5295 | 1.0344 | 0.3864 | 0.9756 | 0.4543 |
| H-MT vs Control | 1.5469 | 0.1021 | 1.2024 | 0.2336 | 0.86243 | 0.5446 | 0.88789 | 0.5383 | 1.3229 | 0.165 | 1.1624 | 0.2566 |
| L-MT vs Control | 1.0831 | 0.3323 | 0.87236 | 0.5665 | 1.1843 | 0.2592 | 0.92217 | 0.511 | 1.7997 | **0.0347*** | 1.2259 | 0.2114 |
| **Soil W** | | | | | | | | | | | | |
| H-MT vs L-MT | 1.3697 | 0.1819 | 1.2114 | 0.2477 | 1.5938 | 0.0713 | 1.3287 | 0.159 | 1.1572 | 0.2749 | 2.1516 | **0.0117*** |
| H-MT vs Control | 2.5237 | **0.0085**** | 1.7198 | 0.0757 | 1.5971 | 0.0879 | 0.9977 | 0.4244 | 1.5354 | 0.0969 | 2.3936 | **0.0092**** |
| L-MT vs Control | 2.0116 | **0.0277*** | 1.0119 | 0.3928 | 1.042 | 0.3649 | 0.8445 | 0.5893 | 0.96995 | 0.4321 | 1.221 | 0.2209 |

H-MT: High melatonin; L-MT: Low melatonin. All treatments and controls for the same soil were composed of a standardised Amount of dilute ethanol. n=4 replicates per treatment. Significance of PERMANOVA (highlighted in bold and shaded): *: 0.01 < p-(MC) ≤ 0.05; **: 0.001 < p-(MC) ≤ 0.01; ***: p-(MC) ≤ 0.001.

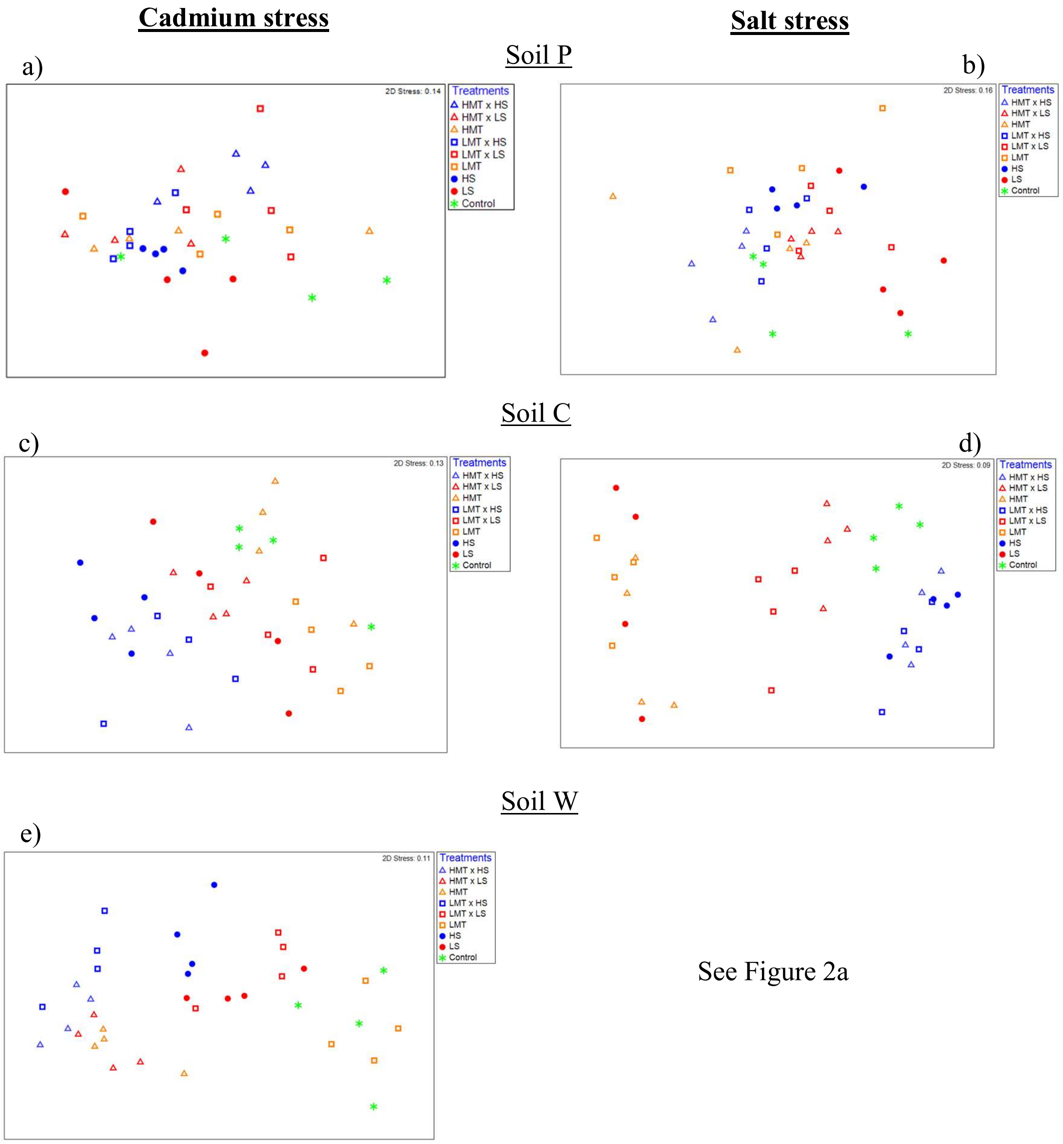


**Supplementary Figure 1**: Non-metric multidimensional scaling (nMDS) ordination displaying Bray-Curtis similarities for bacterial samples within soils P (a & b), C (c & d); and W (e) for various treatments of melatonin and stressor based upon community compositions determined by ARISA fingerprinting analysis. Low 2D stress values indicate high quality ordination plots. Relative proximity of replicates reflects high community similarity within the same treatments for bacterial communities. Water replaced salt treatment and dilute ethanol replaced melatonin treatments in respective control samples. All treatments and controls were composed of a standardised amount of dilute ethanol. (HMT: High melatonin; LMT: Low melatonin; HS: High stress; LS: Low stress; HMT x HS: High melatonin with high stress; HMT x LS: High melatonin with low stress; LMT x HS: low melatonin with high stress; LMT x LS: Low melatonin with low stress).

**Cadmium stress** **Salt stress**

Soil P

a)

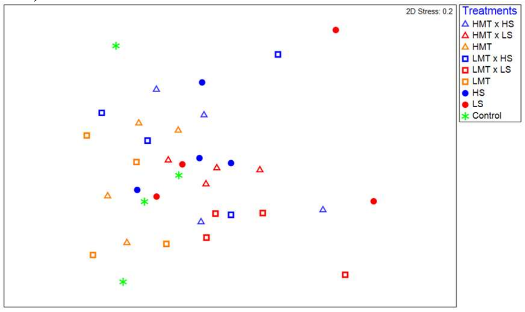


b)

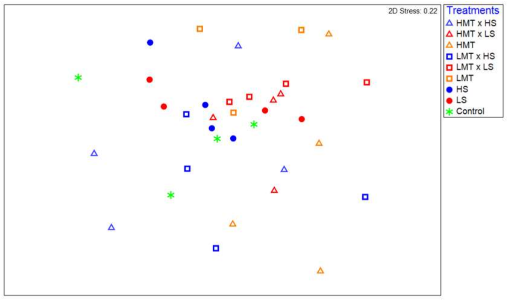


Soil C

c)

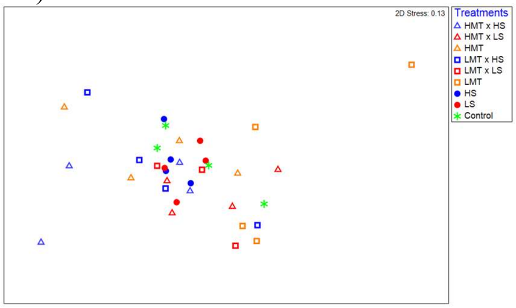


d)

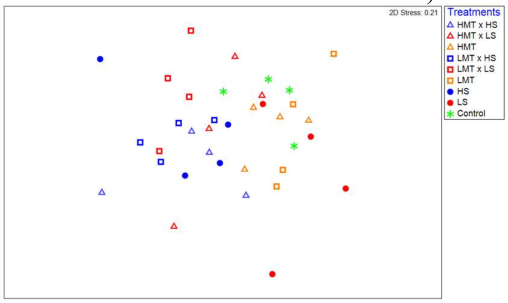


Soil W

e)

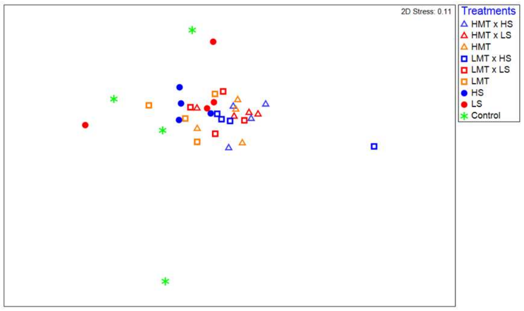


See Figure 3a

**Supplementary Figure 2**: Non-metric multidimensional scaling (nMDS) ordination displaying Bray-Curtis similarities for fungal samples within soils P (a & b), C (c & d); and W (e) for various treatments of melatonin and stressor based upon community compositions determined by ARISA fingerprinting analysis. Low 2D stress values indicate high quality ordination plots. Relative proximity of replicates reflects high community similarity within the same treatments for bacterial communities. Water replaced salt treatment and dilute ethanol replaced melatonin treatments in respective control samples. All treatments and controls were composed of a standardised amount of dilute ethanol. (HMT: High melatonin; LMT: Low melatonin; HS: High stress; LS: Low stress; HMT x HS: High melatonin with high stress; HMT x LS: High melatonin with low stress; LMT x HS: low melatonin with high stress; LMT x LS: Low melatonin with low stress).

#### Supplementary Data 2

##### Melatonin reduced bacterial Shannon species diversity (H') and enhanced fungal Shannon species diversity under low abiotic stress conditions

Shannon species diversity indices (H') calculated the abundance and evenness of OTUs across all soils for bacterial and fungal samples. Non-parametric pairwise Wilcoxon tests were conducted for samples within each soil and stressor treatment to determine significant differences in diversity upon the availability of melatonin. All treatments also contained a standardised dilute concentration of ethanol (approx. 0.05% v/v). Community responses to melatonin under low stress (Supplementary Figure 3) and high stress (Supplementary Figure 4) conditions were analysed.

The average Shannon's diversity index ranged from 2.73 to 3.50 for bacteria and 1.75 to 2.35 for fungi. Bacterial OTU Shannon diversity indices (H') were similar ($p > 0.05$) between control samples (0.05% v/v EtOH) and low stressor only treatments for five of the six experiments, with only low salt stress in soil C resulting in significant differences in bacterial diversity under this comparison ($p < 0.05$) (Supplementary Figure 3a). Diversity decreased in all bacterial communities impacted by melatonin under low stress conditions. Relative to the low stressor treatment within each soil, high melatonin resulted in significant decreases ($p < 0.05$) in bacterial diversity under low cadmium or salt conditions in soil W, as well as in soil C under low salt stress only. In contrast, low melatonin only resulted in a diversity shift ($p < 0.05$) in one soil treatment relative to the low stressor treatment - low salt stress in soil C ($p < 0.05$) (Supplementary Figure 3a).

Fungal communities within control samples (dilute ethanol only) and low stressor (cadmium or salt) treatments showed no significant difference in overall species richness and evenness (H') across all soils for both stressors (Supplementary Figure 3b). In contrast to bacteria, diversity increased in all fungal assemblages impacted by melatonin under low stress conditions. A significant shift in diversity of fungal communities relative to the low stress treatment was only recorded in soil C under low salt stress upon treatment with low melatonin. Communities within soil W under low cadmium stress increased in diversity in response to low melatonin, however only significantly ($p < 0.05$) when compared to the control (Supplementary Figure 3b).

Similar patterns were observed under high stress treatments, with bacterial communities decreasing in diversity in response to melatonin whereas fungi responding to melatonin by increasing in diversity (Supplementary Figure 4). Treatments of melatonin with high cadmium or salt in soil W resulted in significant decreases in bacterial diversity in comparison with the high stressor treatments, whereas only one fungal community (soil W under high cadmium stress) increased significantly ($p < 0.05$) in diversity under the same relative comparison. Otherwise, melatonin showed no effect on microbial community diversity indices under high stress conditions.

a)

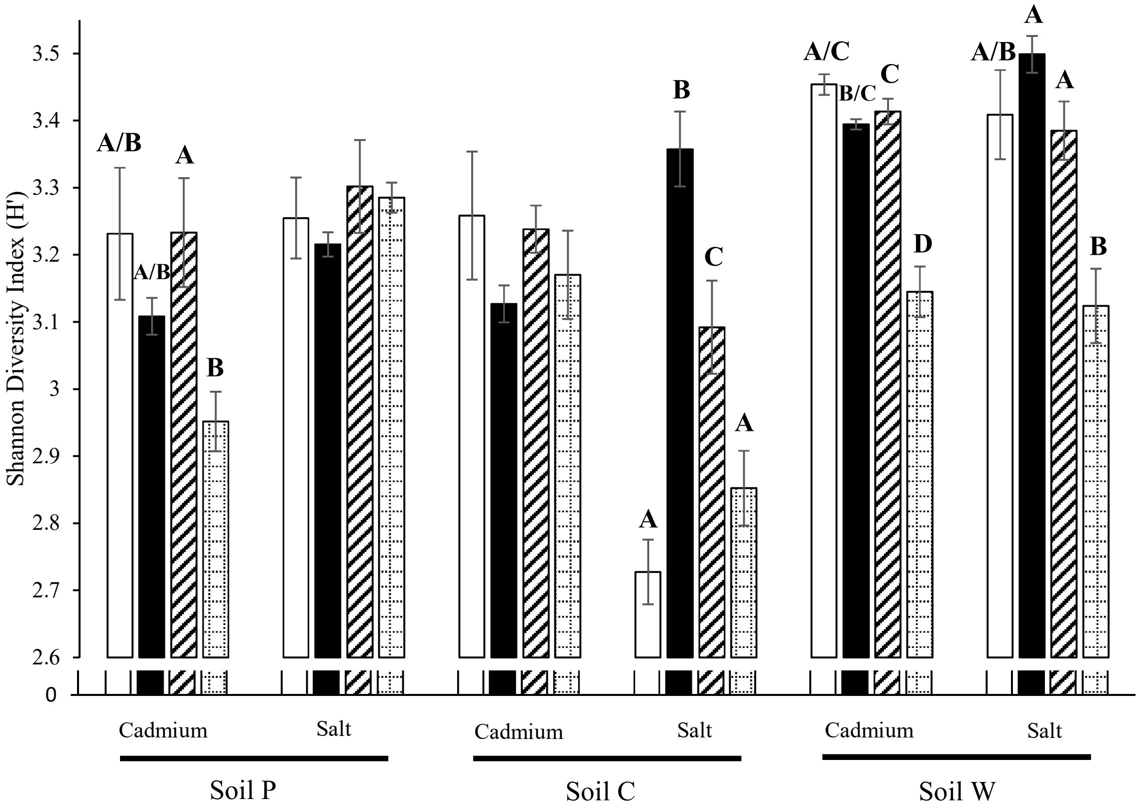

Shannon Diversity Index (H')
3.5
3.4
3.3
3.2
3.1
3
2.9
2.8
2.7
2.6
0
A/B
A/B
A
B
B
C
A
A
A/C
B/C
C
D
A/B
A
A
B
Cadmium
Salt
Cadmium
Salt
Cadmium
Salt
Soil P
Soil C
Soil W

b)

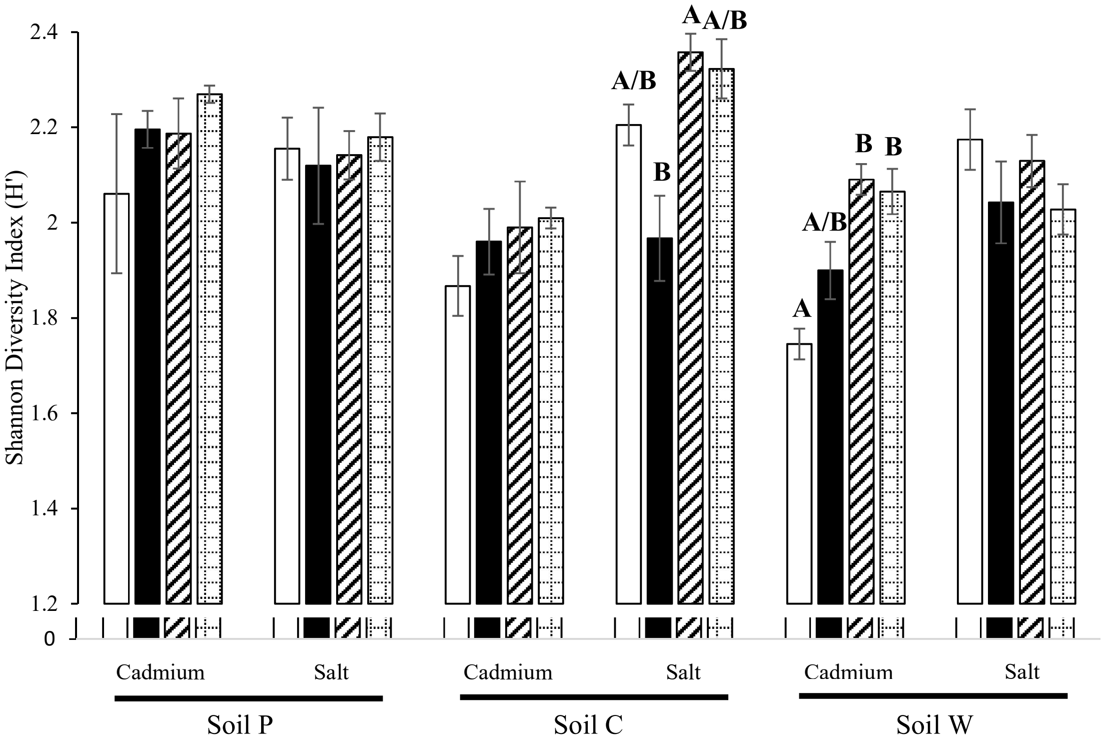


**Supplementary Figure 3.** Mean OTU abundance and evenness based upon OTU diversity (Shannon's index – H') for a) bacterial and b) fungal communities showing the effects of melatonin under low stressor (cadmium or salt) conditions. = Control (no melatonin, no stress); = No melatonin, low stress); = Low melatonin, low stress; = high melatonin, low stress. Shannon's index was calculated in the vegan R software using OTU counts and relative abundances. Bars represent standard error (n = 4). Letters denote significant ($p < 0.05$) differences between treatments within an individual soil experiment as determined by Wilcoxon non-parametric analyses. All treatments and controls were composed of a standardised amount of dilute ethanol per soil.

a)

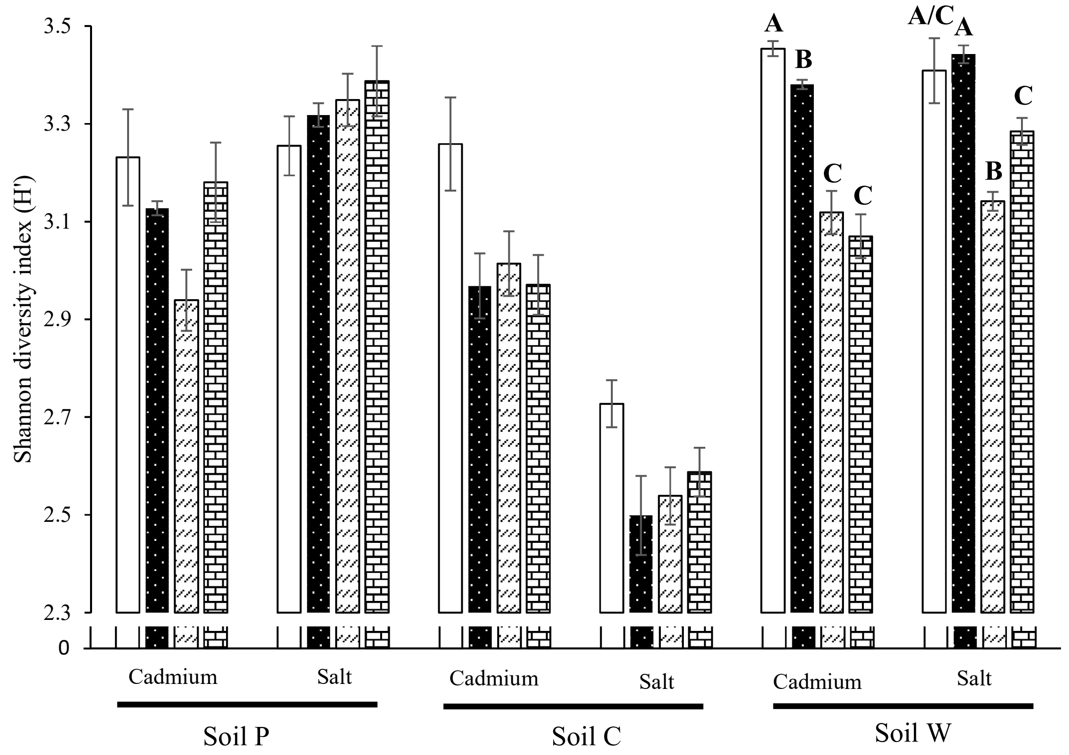

Shannon diversity index (H')
3.5
3.3
3.1
2.9
2.7
2.5
2.3
0
A
B
C
C
A/C
A
B
C
Cadmium
Salt
Cadmium
Salt
Cadmium
Salt
Soil P
Soil C
Soil W


b)

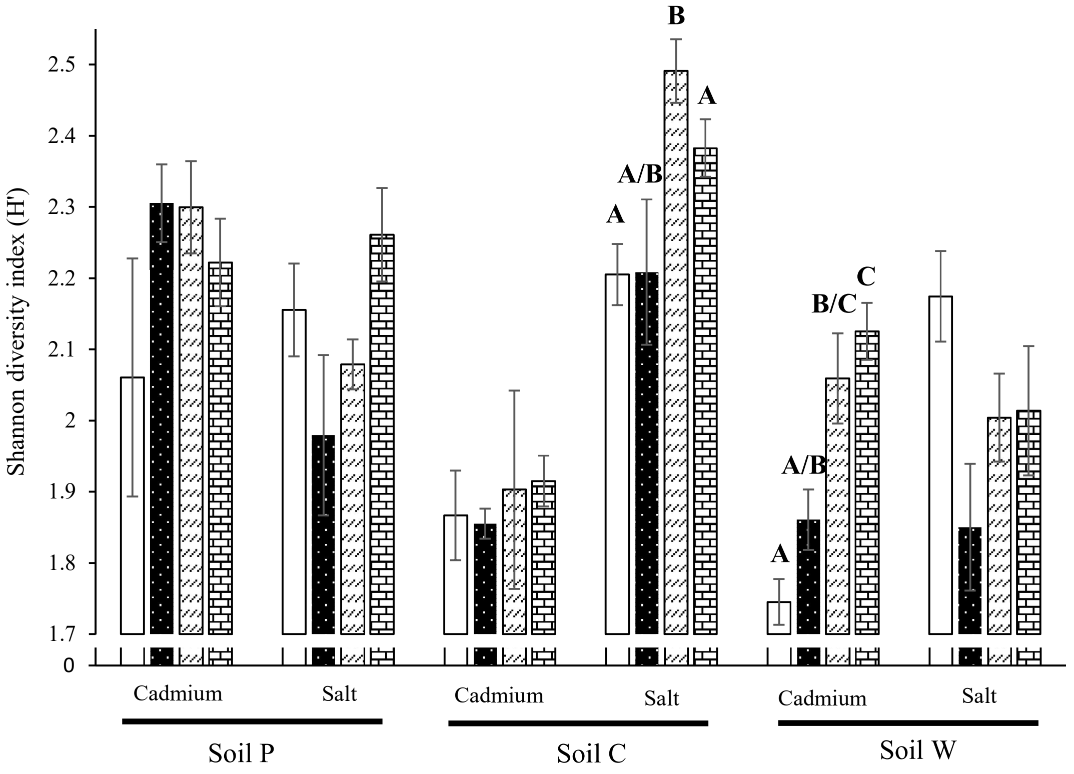

Shannon diversity index (H')
2.5
2.4
2.3
2.2
2.1
2
1.9
1.8
1.7
0
B
A
A/B
A
C
B/C
A/B
A
Cadmium
Salt
Cadmium
Salt
Cadmium
Salt
Soil P
Soil C
Soil W

**Supplementary Figure 4.** Mean OTU abundance and evenness based upon OTU diversity (Shannon's index – H') for a) bacterial and b) fungal communities showing the effects of melatonin under high stressor (cadmium or salt) conditions. = Control (no melatonin, no stress); = No melatonin, low stress; = Low melatonin, low stress; = high melatonin, low stress. Shannon's index was calculated in the vegan R software using OTU counts and relative abundances. Bars represent standard error ($n = 4$). Letters denote significant ($p < 0.05$) differences between treatments within an individual soil experiment as determined by Wilcoxon non-parametric analyses. All treatments and controls were composed of a standardised amount of dilute ethanol per soil.

**Supplemental Data 3**

**Effects of treatments on total bacteria and fungi gene copy numbers for DNA of the 16s rRNA and ITS region respectively**

Total microbial gene copy numbers were assessed to inform about changes in microbial biomass in response to treatments (Supplementary Figures 5 & 6). Soils treated with melatonin show no changes in bacterial biomass compared to control soil samples (i.e. not treated with melatonin or stressors) for soils P and W, whereas high melatonin alone resulted in a significant increase ($p < 0.05$) in soil C, however only at sampling timepoint T0 (Supplementary Figure 5). Total fungal biomass was unaffected by melatonin across all three soils in the absence of an abiotic stressor (Supplementary Figure 6). The effects of melatonin on bacteria and fungi under abiotic stress varied according to stressor concentration and soil type. For example, under low cadmium stress conditions, melatonin application resulted in a significant increase ($p < 0.05$) in total bacteria in soil W, while in soil P, melatonin reduced bacterial gene copy numbers (Supplementary Figure 5a). In contrast, bacteria numbers were unaffected by melatonin at high cadmium concentration. Fungal gene copy numbers also reduced ($p < 0.05$) upon treatment with melatonin under both cadmium concentrations in soil P, however no effects were observed in soils C or W for either stressor condition (Supplementary Figure 6a). Under low salt stress, total bacterial gene copy numbers increased ($p < 0.05$) to the availability of melatonin in soil C, with no shifts observed in the other soil communities under low or high salt stress (Supplementary Figure 5b). Total fungal gene copy numbers decreased in soils P and W under salt stress upon the availability of melatonin, however no differences were observed in soil C. (Supplementary Figure 6b).

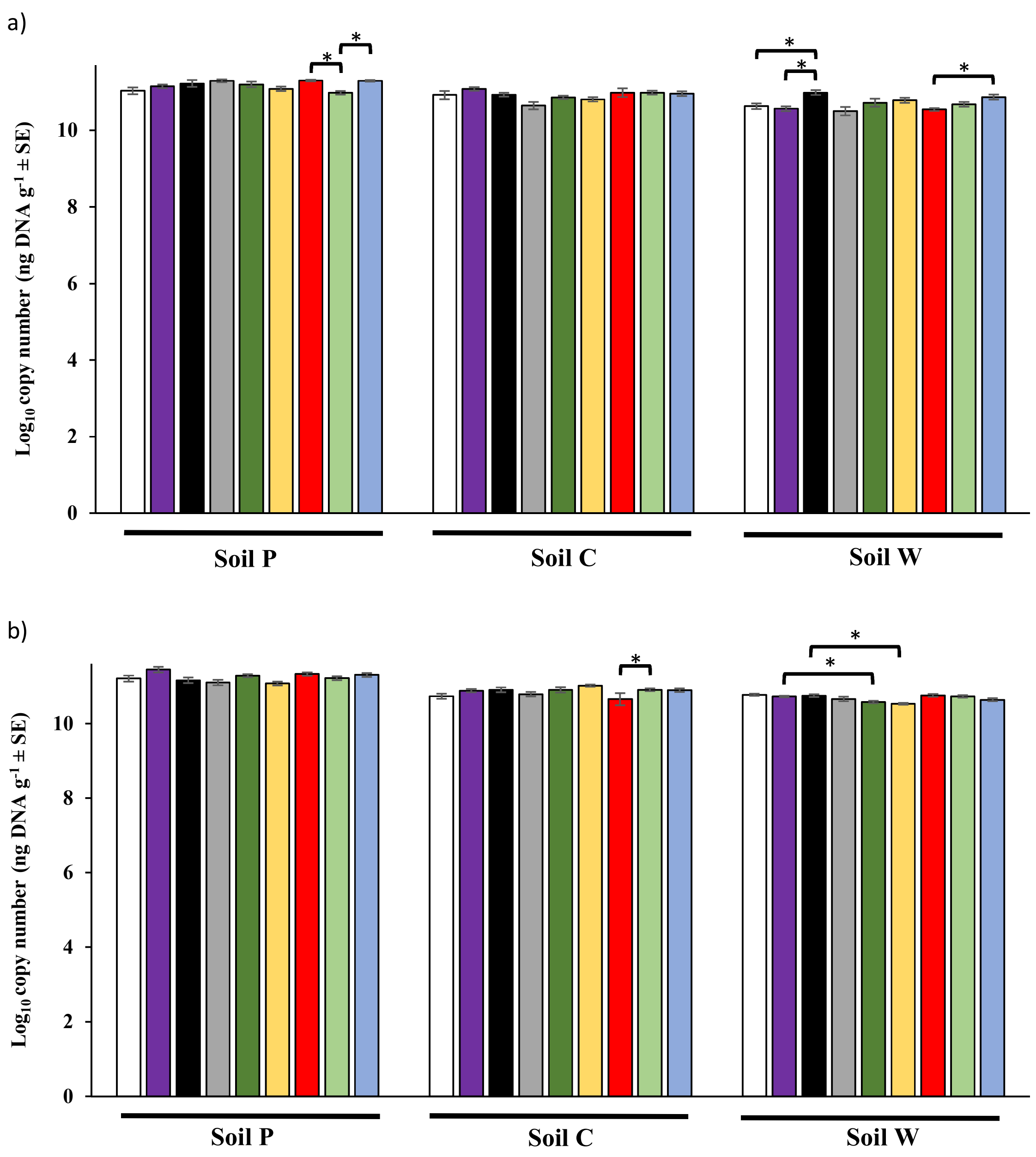


**Supplementary Figure 5**: Quantitative PCR estimation of bacterial (16S rRNA) gene copy copy numbers for communities treated with a) cadmium and b) salt stressors. Asterisks (*) represents significant differences ($p < 0.05$) in copy number for communities treated with melatonin-only compared to control samples or high / low stressor (cadmium or salt) in the presence vs absence of melatonin as determined by pairwise Wilcoxon non-parametric analyses. Treatments: = Control; = low melatonin; = high melatonin; = high stress; = low melatonin + high stress; = high melatonin + high stress; = low stress;

= low melatonin + low stress; = high melatonin + low stress. Values (± SE) are reported as mean of n = 4 biological replicates based on an average weight of 0.275g soil per sample. All treatments and controls were exposed to a standard amount of dilute ethanol per soil.

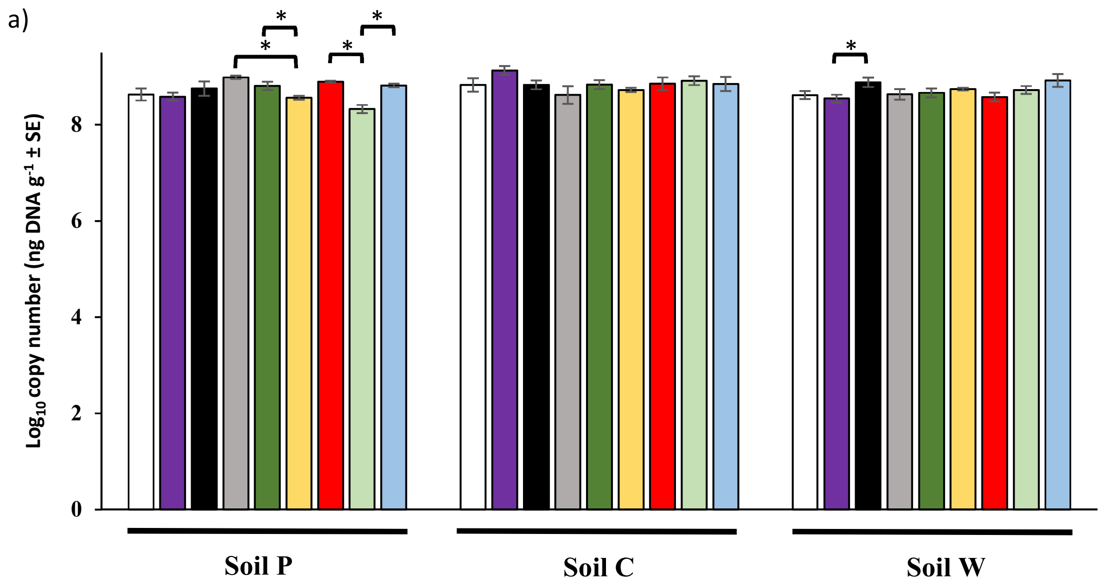


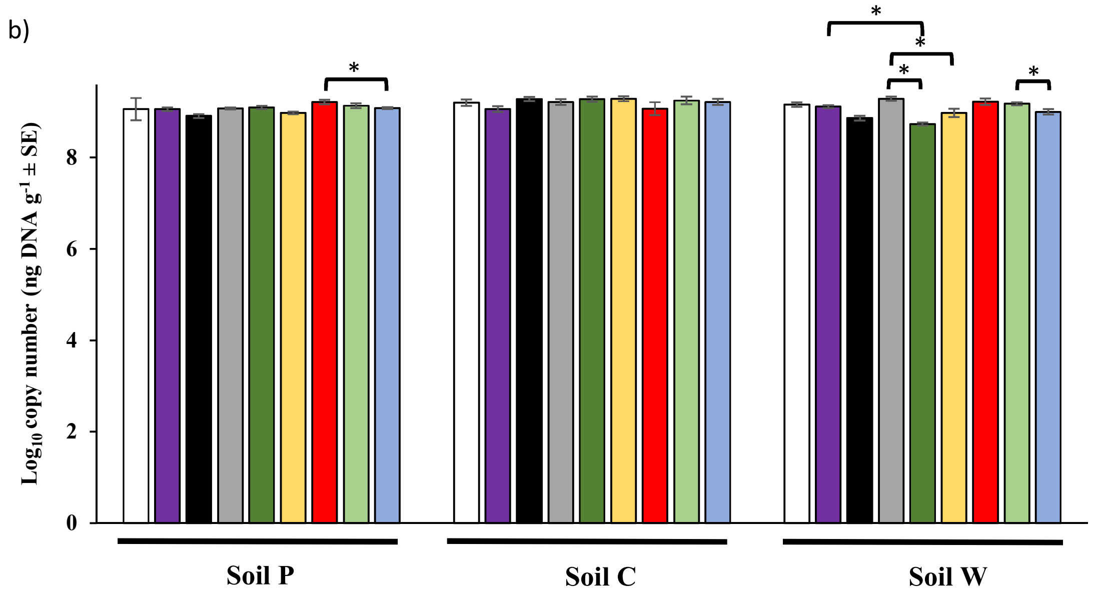


**Supplementary Figure 6:** Quantitative PCR estimation of fungal (ITS) gene copy numbers for communities treated with a) cadmium and b) salt stressors. Asterisks (*) represents significant differences ($p < 0.05$) in copy number for communities treated with melatonin-only compared to control samples or high / low stressor (cadmium or salt) in the presence vs absence of melatonin as determined by pairwise Wilcoxon non-parametric analyses. Treatments: □ = Control; ■ = low melatonin; ■ = high melatonin; ■ = high stress; ■ = low melatonin+high stress; ■ = high melatonin+high stress; ■ = low stress; ■ = low melatonin+low stress; ■ = high melatonin+low stress. Values (± SE) are reported as mean of n = 4 biological replicates based on an average weight of 0.275g soil per sample. All treatments and controls were exposed to a standard amount of dilute ethanol per soil.

**Supplementary Data 4**

**Melatonin and/or stressor treatments altered microbial community similarities**

Analysis of dissimilarity of microbial community structures between control and treatment (melatonin and/or stressor) soil samples showed varying trends for both bacteria and fungi (Ranjard et al., 2001, Chow et al., 2013) (Supplementary Table 7). Under low salt conditions, melatonin-treated bacterial samples (green highlight) showed a dramatic decrease in dissimilarity to control compared to low salt only (yellow highlight) samples across all three soils (P: 7.6%; C: 33.5%; W: 19.7%) (Supplementary Table 7a). However, this trend was not observed at high salt conditions for bacterial samples, as melatonin had little effect on the relative differences in dissimilarity, in some cases further increasing dissimilarity. Under high cadmium treatments, melatonin further increased dissimilarity to control compared to the high cadmium-only samples for bacteria, with variable results observed under low cadmium conditions. For fungal communities, changes in dissimilarity under abiotic stress conditions did not follow this same trend at all, with varying patterns across the three soils.

**Supplementary Table 7**: Dissimilarities (%) of a) bacterial and b) fungal communities treated with melatonin and/or stressor in comparison to community structures within control samples as determined by SIMPER using Bray Curtis dissimilarity analysis. Colour codes indicate changes in dissimilarities in bacterial communities under low salt stress conditions between samples treated with high melatonin (green) in comparison to no melatonin (yellow).

**a)**

| | **Soil P** | | | **Soil C** | | | **Soil W** | | |
|---|---|---|---|---|---|---|---|---|---|
| | **High MT** | **Low MT** | **No MT** | **High MT** | **Low MT** | **No MT** | **High MT** | **Low MT** | **No MT** |
| **High Cd** | 42.53 | 40.78 | 37.98 | 41.16 | 40.17 | 40.97 | 53.83 | 52.97 | 41.22 |
| **Low Cd** | 43.45 | 40.83 | 44.3 | 29.15 | 31.75 | 34.02 | 49.35 | 34.44 | 34.42 |
| **No Cd** | 41.3 | 39.89 | - | 27.13 | 31.76 | - | 46.58 | 28.1 | - |
| **High salt** | 36.28 | 32.84 | 35.87 | 35.63 | 36.54 | 31.81 | 41.25 | 41.75 | 37.02 |
| **Low salt** | 32.54 | 36.97 | 40.17 | 29.45 | 43.62 | 62.98 | 34.14 | 24.98 | 44.67 |
| **No salt** | 37.57 | 40.57 | - | 61.71 | 61.58 | - | 50.46 | 48.05 | - |

**b)**

| | Soil P | | | Soil C | | | Soil W | | |
|---|---|---|---|---|---|---|---|---|---|
| | **High MT** | **Low MT** | **No MT** | **High MT** | **Low MT** | **No MT** | **High MT** | **Low MT** | **No MT** |
| **High Cd** | 27.94 | 29.64 | 26.74 | 27.16 | 24.44 | 18.76 | 35.59 | 39.05 | 27.81 |
| **Low Cd** | 26.87 | 31.33 | 31.8 | 21.74 | 20.1 | 19.85 | 34.8 | 31.54 | 30.32 |
| **No Cd** | 26.24 | 25.16 | | 23.77 | 28.84 | | 33.72 | 28.54 | |
| **High salt** | 24.34 | 22.88 | 19.55 | 26.85 | 25.96 | 26.21 | 29.55 | 29.79 | 32.16 |
| **Low salt** | 21.01 | 21.56 | 21.4 | 22.2 | 26.97 | 26.74 | 26.66 | 21.73 | 20.64 |
| **No salt** | 25.09 | 21.95 | | 20.29 | 21.71 | | 26.24 | 18.2 | |

MT: melatonin; Cd: cadmium. All treatments and controls were composed of a standardised amount of dilute ethanol per soil. n=4 replicates per treatment.

a)

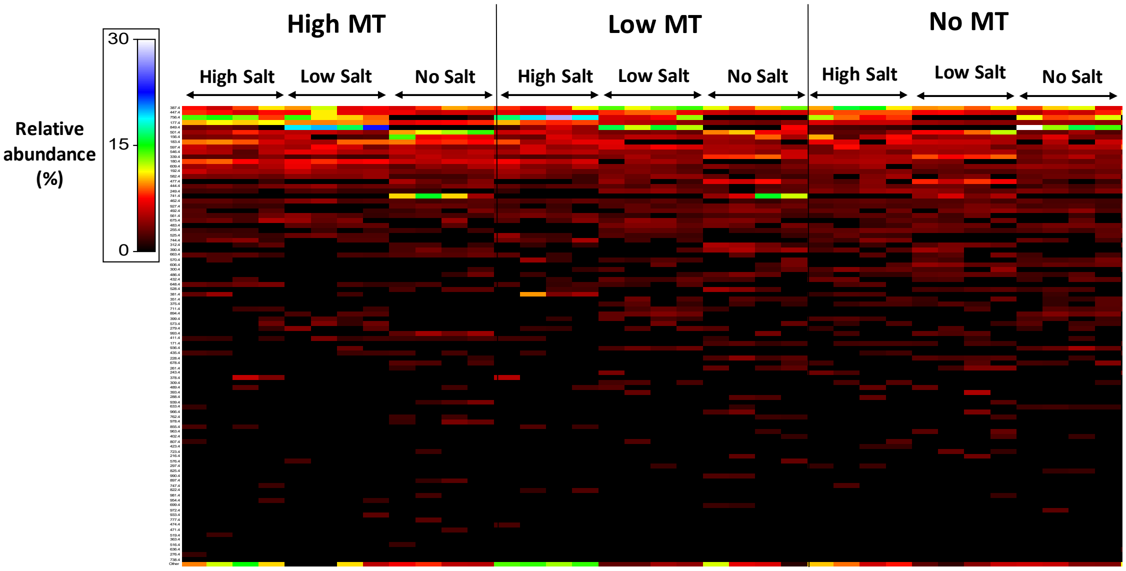


b)

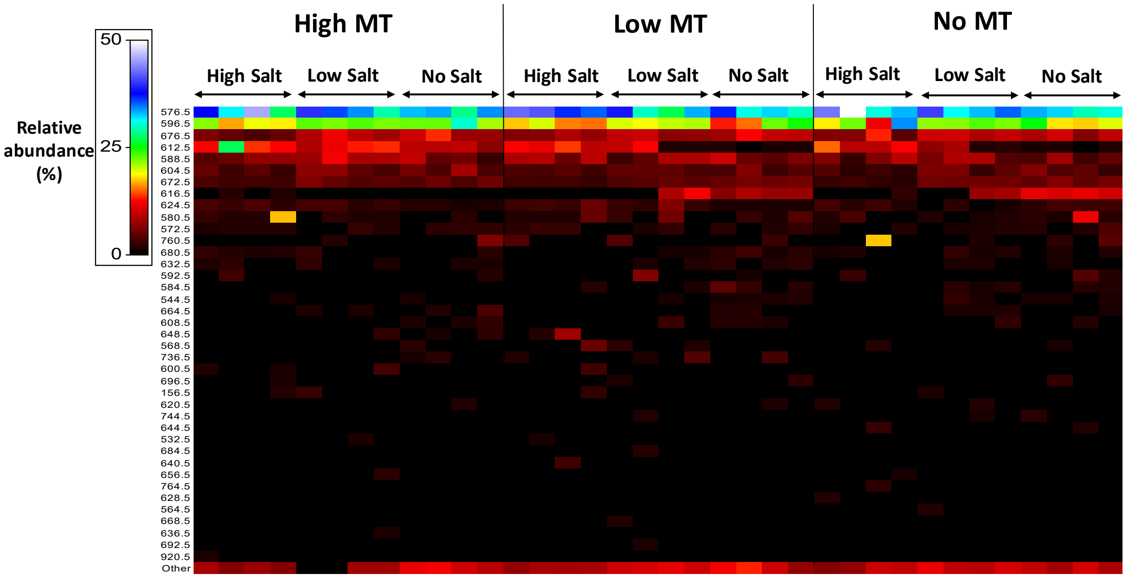


**Supplementary Figure 7:** Heatmaps of a) bacterial and b) fungal community compositions for treatments with melatonin and salt in soil W. OTU identifications are provided in rows and columns represents replicate samples (n=4) for various treatments. Relative abundances (%) of individual OTUs are indicated by a colorimetric scale ranging from 0-30% for bacterial and 0-50% for fungal samples. Singletons and OTUs less than 1% relative abundance were removed and classified within the category ‘Other’, listed as the final row. MT: melatonin. All treatments and controls were composed of a standardised amount of dilute ethanol.

#### References


Chow, C. E. T., Sachdeva, R., Cram, J. A., Steele, J. A., Needham, D. M., Patel, A., Parada, A. E. and Fuhrman, J. A. (2013). Temporal variability and coherence of euphotic zone bacterial communities over a decade in the Southern California Bight. *ISME Journal* 7, 2259-2273. doi: 10.1038/ismej.2013.122.

Hansgate, A. M., Schloss, P. D., Hay, A. G. and Walker, L. P. (2005). Molecular characterization of fungal community dynamics in the initial stages of composting. *FEMS Microbiology Ecology* 51, 209-214. doi: 10.1016/j.femsec.2004.08.009.

Ranjard, L., Poly, F., Lata, J. C., Mougel, C., Thioulouse, J. and Nazaret, S. (2001). Characterization of Bacterial and Fungal Soil Communities by Automated Ribosomal Intergenic Spacer Analysis Fingerprints: Biological and Methodological Variability. *Applied and Environmental Microbiology* 67, 4479-4487. doi: 10.1128/AEM.67.10.4479-4487.2001.
